# Reversible One-way Lipid Transfer at ER-Autophagosome Membrane Contact Sites via Atg2

**DOI:** 10.1101/2025.03.27.645728

**Authors:** Li Hao, Tomoki Midorikawa, Yuta Ogasawara, Takuma Tsuji, Takuma Kishimoto, Yutaro Hama, Huichao Lang, Nobuo N. Noda, Kuninori Suzuki

## Abstract

Bridge-like lipid transfer proteins (LTPs) contain a repeating β-groove domain and long hydrophobic grooves that act as bridges at membrane contact sites (MCSs) to efficiently transfer lipids. Atg2 is one such bridge-like LTP essential for autophagosome formation, during which a newly synthesized isolation membrane (IM) emerges and expands through lipid supply. However, studies on Atg2-mediated lipid transfer are limited to *in vitro* studies due to the lack of a suitable probe for monitoring phospholipid dynamics *in vivo*. Here, we characterized the lipophilic dye octadecyl rhodamine B (R18), which internalizes and labels the endoplasmic reticulum (ER) in a manner that requires flippases and oxysterol-binding protein–related proteins. Using R18, we demonstrated phospholipid transfer from the ER to the IM during autophagy *in vivo*. Upon autophagy termination, we observed reversible phospholipid flow from the IM to the ER in response to environmental changes. Our findings highlight the critical role of bridge-like LTPs in MCS-mediated phospholipid homeostasis.

## Introduction

Phospholipids are key constituents of various organelle membranes in eukaryotic cells. They are regulated asymmetrically and heterogeneously to form permeability barriers that separate organelles from their surroundings, creating distinct compartments. This organization facilitates the segregation and regulation of numerous reactions within cells. Cells maintain lipid homeostasis in organelle membranes by actively translocating and transferring lipids, ensuring optimal membrane function.

Membranes exchange phospholipids with each other via vesicular and nonvesicular pathways. In vesicular trafficking, vesicles transport lipids through secretory and endocytic pathways^1^. However, this process has several limitations. First, the quantity and rate at which vesicles deliver lipids are insufficient for organelle biogenesis^2^. Second, lipid transfer occurs in organelles that do not receive vesicles^3^. Therefore, it is plausible that pathways involved in the transport of phospholipids via mechanisms other than vesicular transport are crucial for maintaining cellular functions^4–7^. Nonvesicular trafficking primarily relies on lipid transfer proteins (LTPs) at membrane contact sites (MCSs), where the two organelles physically interact. Lipid transfer at these sites typically affects the functions of either organelle alone or both organelles^8–10^. The most well-characterized LTPs have a box-like structure with a hydrophobic core, allowing them to hold one or a few lipids^10–12^. These proteins transport lipids by first docking onto the donor membrane, extracting specific lipids. They diffuse through the cytoplasm before inserting the lipids into the acceptor membrane^10, 13^. Recent studies have identified a new class of bridge-like LTPs characterized by repeating β-groove (RBG) domains and extended hydrophobic grooves^14, 15^. These large structures allow LTPs to bridge MCSs and facilitate efficient lipid transfer^7^. Five RBG proteins—Vps13, Atg2, BLTP1 (Tweek/Csf1/KIAA1109), BLTP2 (Hob/Fmp27/KIAA0100), and BLTP3 (SHIP164)^15–24^—have been identified in eukaryotes.

Among the bridge-like LTPs, Atg2 plays a critical role in macroautophagy (herein referred to as autophagy) while Vps13 has been proposed to exhibit a parallel function^25–27^. Autophagy is a universally conserved intracellular degradation pathway that plays an essential role in cell homeostasis and stress responses^28^. It relies on the reorganization of lipid membranes to form an isolation membrane (IM), also known as the phagophore, between the endoplasmic reticulum (ER) and vacuole, which contains engulfed cytoplasmic components. The IM then closes to become an autophagosome, enclosing damaged organelles or protein aggregates for degradation in vacuoles or lysosomes^29–31^. A substantial number of phospholipids is transported to support autophagosome biogenesis, and recent studies have further highlighted the crucial roles of *de novo* phospholipid synthesis in facilitating IM expansion^32–34^. Atg2 forms a complex with Atg18 and the scramblase Atg9 at the edge of the expanding IM during autophagosome formation^35–38^. Atg2-Atg18 complex also constricts IM rims in a phosphatidylinositol 3-phosphate (PI(3)P) dependent manner, promoting autophagosome closure^39^. Additionally, ER exit sites (ERES) have been suggested as a key hub for Atg2 localization during autophagosome biogenesis^40, 41^. Previous *in vitro* studies on the crystal structure of Atg2 N-terminus suggest that it directly participates in phospholipid transfer at the ER–IM MCS^18^. However, the organelles responsible for the lipid supply for IM expansion are uncertain^42–46^.

Considerable progress has recently been made in our understanding of nonvesicular lipid transport. Most evidence supporting the Atg2-mediated lipid transfer model is largely based on *in vitro* studies^17, 18, 25, 47, 48^. However, significant gaps remain in our understanding of direct lipid transfer in living cells. The lack of tools for determining lipid transfer in living cells raises concerns about the validity of conclusions drawn from such studies in a real cellular environment. Additionally, the directionality of lipid transfer by Atg2 remains unclear *in vivo*.

In this study, to investigate lipid transfer during autophagosome formation, we first identified the lipophilic dye octadecyl rhodamine B (R18), which undergoes nonvesicular transport from the plasma membrane (PM) through the ER to autophagy-related structures (ARSs), including the pre-autophagosomal structure/phagophore assembly site (PAS), IMs, autophagosomes, and autophagic bodies. Given that special intracellular transfer route of R18, we investigated lipid transfer from the ER to the IM during autophagosome formation. Our time-lapse imaging analysis also revealed rapid lipid supply accompanied by IM expansion. Our study demonstrated lipid transfer from the ER to the IM and strongly supported the hypothesis that the ER is the primary source of lipids for IM expansion. Furthermore, we developed a repletion assay using a nutrient-rich medium to terminate autophagy under the fluorescence microscope. We identified reversed lipid transfer from the IM to the ER by measuring the recovery of R18 fluorescence at the ER–IM MCS after photobleaching. These findings indicate a reversible one-way lipid transfer mediated by Atg2 at the ER–IM MCS in response to environmental changes. Our research provides direct *in vivo* evidence of the reversible lipid transfer mediated by the bridge-like LTP Atg2, offering new insights into the dynamic regulation of lipid homeostasis during autophagy.

## Results

### R18 stains the ER and each autophagy-related membrane structure

R18 is a lipophilic fluorescent dye with an alkyl tail composed of 18 carbon atoms, allowing it to be easily inserted into phospholipid membranes. In the last century, it was used to stain membranes in *in vitro* assays; however, its use was gradually phased out with the development of a series of fluorescently labeled phospholipids^49, 50^. We assessed the ability of R18 to label ARSs by performing colocalization analysis using yeast cells expressing Atg8, a marker of ARSs, tagged with mNeonGreen (mNG). mNG–Atg8 was visualized as a distinct dot corresponding to the PAS in wild-type (WT) cells (Figure 1A and 1B). In contrast, R18 signal was not recognized at the mNG-Atg8 labeled dots in *atg2*Δ cells (Figure 1A and 1B). The Atg2–Atg18 complex is absent in the PAS of *atg2*Δ cells^38^. Besides, R18 accumulates at the PAS in Atg8^G116^ *atg4*Δ cells, where the IM fails to expand but Atg2 is still recruited^51^. Taken together, these results suggest that the Atg2–Atg18 complex is involved in the staining of the PAS with R18. Because the IM appears as a dot in WT cells, we enlarged it into a cup-shaped structure by overexpressing prApe1, a selective autophagy cargo (see Figure 4B)^40^. We found that mNG–Atg8 labeled a cup-shaped IM and colocalized with R18 (Figure 1A), as previously reported^51^. We then used the W303 background cells to examine whether a ring-shaped autophagosome could be stained with R18^41^. We found that R18 was colocalized with ring-shaped autophagosomes (Figure 1A and 1B). These data collectively demonstrate that R18 is a fluorescent dye capable of *in vivo* labeling of IMs and autophagosomes.

**Figure 1.**
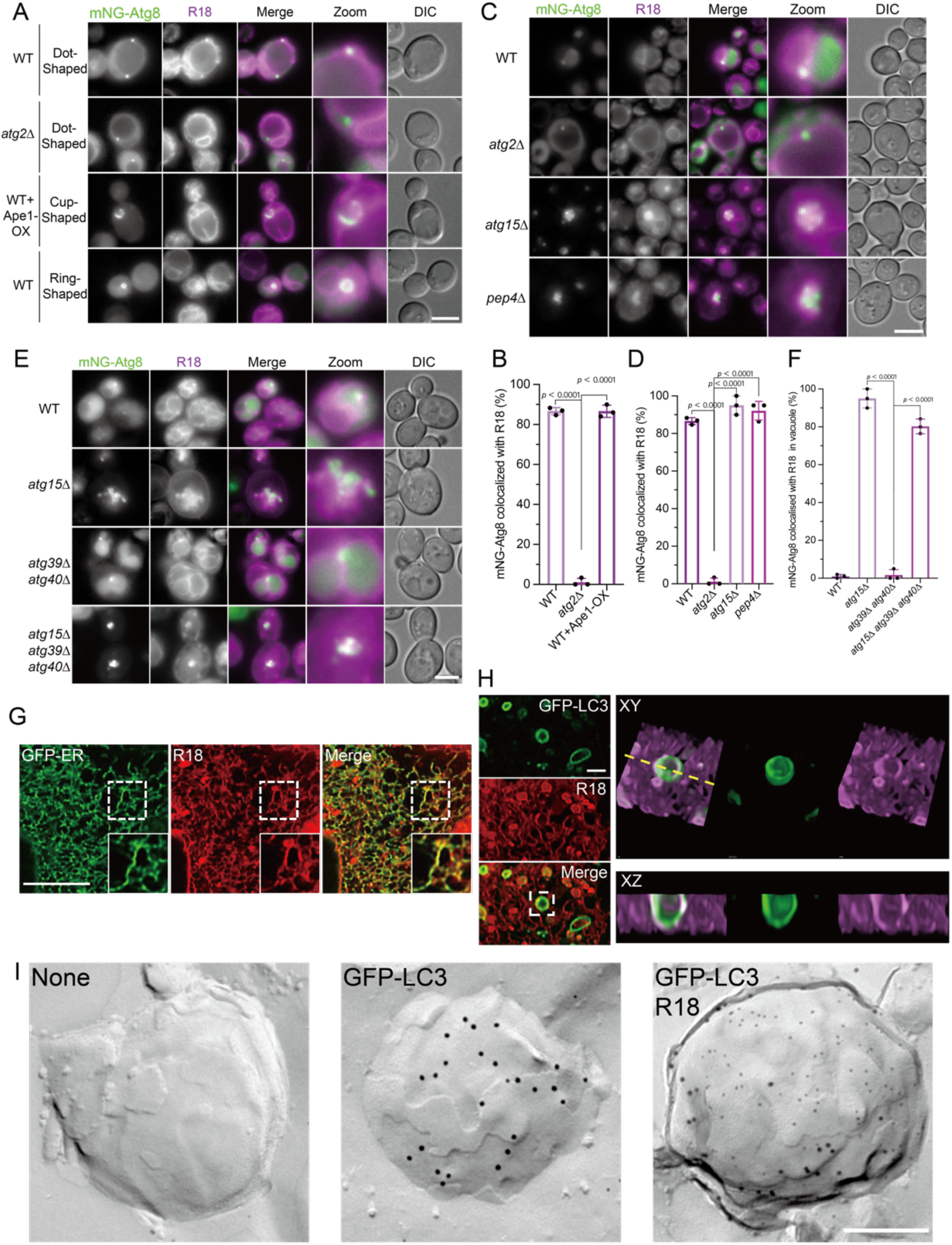
Octadecyl rhodamine B (R18) stains autophagy-related membrane structures. (A) Colocalization between mNG–Atg8 and R18 in W303 background cells treated with rapamycin to induce autophagy. Cells were stained with R18 for 10 minutes, washed with fresh medium, and then treated with 200 ng/mL rapamycin for 1 hour before imaging. The scale bar represents 5 μm. WT, wild-type. OX, overexpression. mNG, mNeonGreen. DIC, differential interference contrast. (B) Percentage of colocalization calculated using 100 cells from 3 independent experiments in (A). WT includes both dot-and ring-like structures of mNG–Atg8 colocalized with R18. Error bars represent standard deviation (SD). P-values were calculated using one-way ANOVA. (C) Colocalization between mNG–Atg8 and R18 in SEY6210 background cells. Cells were stained with R18 for 10 minutes, washed with fresh medium, and then treated with 200 ng/mL rapamycin for 1 hour before imaging. The scale bar represents 5 μm. (D) Percentage of colocalization calculated using 100 cells from 3 independent experiments in (C). Error bars represent SD. P-values were calculated using one-way ANOVA. (E) Colocalization between mNG–Atg8 and R18 in SEY6210 background cells. Cells were stained with R18 for 10 minutes, washed with fresh medium, and then treated with 200 ng/mL rapamycin for 1 hour before imaging. The scale bar represents 5 μm. (F) Percentage of colocalization calculated using 100 cells from 3 independent experiments in (E). Error bars represent SD. P-values were calculated using one-way ANOVA. (G) MEFs expressing GFP–ER were stained with R18 for colocalization analysis. The boxed area represents a higher magnification. (H) MEFs expressing GFP–LC3 were subjected to autophagy induction by starvation and stained with R18 for colocalization analysis. The boxed area represents a higher magnification. (I) Freeze-fracture replica double labeling of R18 and GFP–LC3 in MEF cells. Starved MEFs expressing GFP–LC3 were processed for freeze-fracture replica immunogold labeling. R18 was detected using 6-nm colloidal gold, and GFP–LC3 with 12-nm gold. The scale bar represents 200 nm.

The outer membrane of the autophagosome fuses with the vacuolar membrane. The inner membrane compartment, referred to as the autophagic body, is then released into the vacuolar lumen and degraded^52^. Here, we examined whether the autophagic body was labeled with R18. We found that mNG–Atg8 diffused throughout the vacuolar lumen of WT cells, representing the disintegration of autophagic bodies (Figure 1C and 1D). Conversely, the vacuolar lumen was not stained with mNG–Atg8 in *atg2*Δ cells (Figure 1C and 1D). Atg15 and Pep4 are crucial vacuolar hydrolases for the breakdown of autophagic bodies^53–55^. In *atg15*Δ and *pep4*Δ cells, the mNG–Atg8-labeled autophagic bodies were found to accumulate inside the vacuole and colocalized with R18, indicating that R18 can label autophagic bodies (Figure 1C and 1D). We also combined *atg15*Δ or *pep4*Δ with *atg1*Δ, which completely blocks autophagosome formation. As expected, mNG–Atg8-labeled autophagic bodies did not accumulate inside the vacuole in these double mutants. R18 showed no vacuolar accumulation (Fig. S1A). These results demonstrate that R18 does not enter the vacuole independently of autophagic bodies delivery.

In Figure 1C, we pre-stained cells with R18 and then induced autophagy by rapamycin treatment, resulting that R18 localized to the ER was transferred to autophagic bodies. We then first induced autophagy by rapamycin treatment and then stained cells with R18. Although mNG–Atg8–positive autophagic bodies accumulated in the vacuole, R18 staining of the autophagic bodies was not recognized (Fig. S1B). These findings suggest that R18 cannot stain autophagic bodies by diffusion but by autophagy.

ER-phagy is a type of selective autophagy in which the ER is specifically sequestered within autophagosomes and then degraded^56–58^. Because R18 also stains the ER, the dots visualized with R18 could be R18-labeled ER engulfed into autophagic bodies. We used ER-phagy-deficient cells to exclude this possibility. Atg39 and Atg40 localize to the ER membrane, where they participate in the ER-phagy-dependent degradation of the perinuclear and cortical ER, respectively^58^. The *atg39*Δ *atg40*Δ cells exhibited diffusion of mNG within the vacuole (Figure 1E and 1F), indicating that autophagosomes lacking the ER were transported into the vacuole. In *atg15*Δ or *atg15*Δ *atg39*Δ *atg40*Δ cells, we observed the accumulation of mNG-Atg8-labeled autophagic bodies inside the vacuole (Figure 1E and 1F), suggesting the inhibition of autophagic body degradation. Consistently, we found an accumulation of R18-positive autophagic bodies in these cells (Figure 1E and 1F). To further ensure that R18 puncta were not derived from any selective autophagy pathway, we also analyzed *atg11*Δ *atg39*Δ *atg40*Δ cells, in which all forms of selective autophagy are blocked. In this background, mNG–Atg8 labeled puncta still colocalized with R18, confirming that R18 labels the limiting membrane of autophagic bodies independently of selective autophagy (Figure S1C). Together, these results suggest that the limiting membranes of autophagic bodies are labeled with R18. We conclude that R18 can serve as a traceable marker for the membrane dynamics of each ARS, including the PAS, IMs, autophagosomes, and autophagic bodies.

We further validated the localization of R18 by examining its colocalization with green fluorescent protein (GFP)-tagged organelle marker proteins (Figure S2A). We first used FM4-64, which can stain each compartment along the endocytosis pathway and localize to the vacuolar membrane, to validate our method (Figure S2B and S2C). Subsequently, cells expressing GFP-tagged organelle marker protein-expressing cells were stained with R18. We found that R18 colocalized with the ER membrane marker proteins Dpm1-GFP, Erd2-GFP, and Sec61-GFP, which have distinct subcellular localizations (Figure S2D and S2E). R18 colocalized with these ER markers (Figure S2D and S2E). R18 occasionally labeled the vacuolar membrane, possibly due to the fusion of autophagosomes with the vacuole. However, R18 did not colocalize with other organelle markers (Figure S2D and S2E), indicating that it primarily stains the ER.

Finally, we confirmed whether R18 can be used in mammalian cells by testing it in mouse embryonic fibroblasts (MEFs) expressing GFP-tagged ER marker proteins. Under normal conditions, GFP and R18 exhibited clear colocalization following R18 staining (Figure 1G). After starvation-induced autophagy, we found that R18 colocalized with GFP-LC3, an autophagosome marker (Figure 1H). Cross-sectional imaging confirmed that R18 colocalized with GFP-LC3 in the XY and XZ planes, indicating the suitability of R18 for studying autophagy in mammalian cells (Figure 1H). In mammalian cells, the ER is closely associated with the IM. Thus, it is difficult to distinguish the IM from the ER using fluorescence microscopy. To directly determine whether R18 integrates into autophagosomal membranes, we used freeze-fracture replica electron microscopy, a high-resolution method that preserves membrane phospholipids in a metal–carbon replica^59^. Using this approach, we performed double immunogold labeling of R18 and GFP–LC3. Small (6 nm) gold particles marking R18 and larger (12 nm) particles marking GFP–LC3 were both detected on the autophagosomal membrane, demonstrating that R18 indeed integrates into autophagosomal membranes in starved MEFs (Figure 1I).

### Internalization of R18 depends on flippases at the plasma membrane

To further validate R18 as a reliable probe for tracing biologically regulated lipid transfer — rather than passive diffusion — we next examined its intracellular entry pathway. First, we investigated how R18 enters cells and stains the ER, comparing it to the internalization pathway of the well-established lipophilic dye FM4-64^60^. In contrast to FM4-64, which localizes to endocytic compartments, R18 preferentially stains the ER and ARSs (Figures. 1 and S2). These results suggest that FM4-64 and R18 are internalized and transported via different pathways. FM4-64 is internalized via endocytosis, which depends on physiological temperature and ATP^60^. Here, we examined whether biological processes also influence the internalization of R18. We expressed GFP-tagged Dpm1 as a marker of ER transmembrane proteins. At 30°C, cells displayed clear colocalization of R18 with Dpm1-GFP, which preferentially stained the ER (Figure 2A and 2B). However, at 4°C, a drastic accumulation of R18 on the PM was observed (Figure 2A and 2B), suggesting that the internalization of R18 is temperature-dependent. Next, ATP depletion was achieved by inhibiting ATP synthesis through oxidative respiration and glycolysis using NaN_3_ and NaF, respectively^61^. Without these inhibitors, R18 colocalized with the ER containing Dpm1-GFP (Figure 2C–2E). However, in the presence of these inhibitors, R18 accumulated on the PM, forming a distinct circular pattern (Figure 2D and 2E). We then washed out NaN_3_ and NaF using fresh medium (Figure 2C) and found that R18 rapidly entered the cells, colocalizing with Dpm1-GFP within 3 min (Figure 2D and 2E). This observation indicates the rapid transfer of R18 from the outside of cells to the ER. These results demonstrate that R18 internalization depends on physiological temperature and ATP.

**Figure 2.**
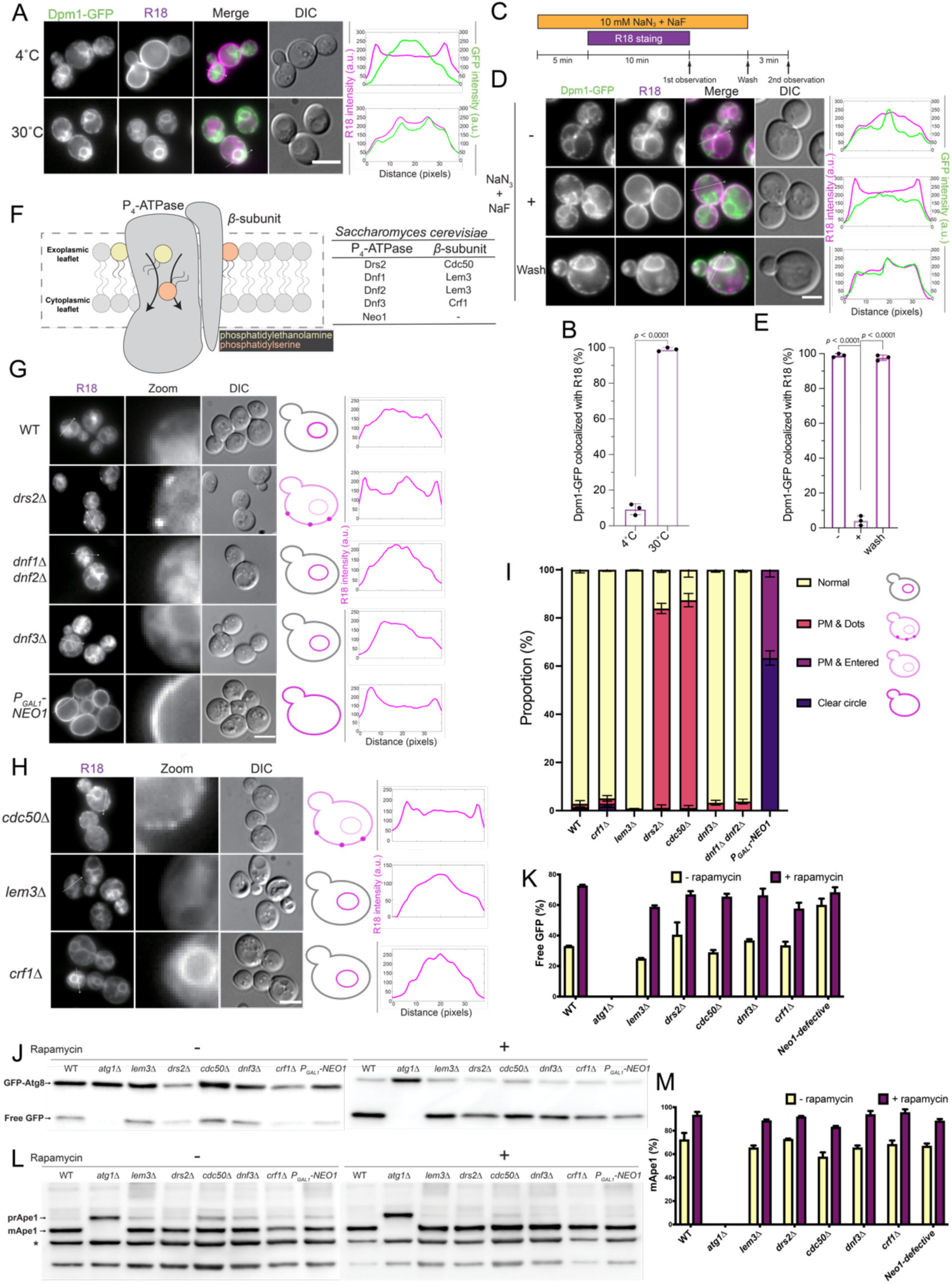
R18 internalization at the plasma membrane requires flippase activity. (A) Colocalization between Dpm1–GFP and R18 in WT cells. The cells were stained with R18 at the indicated temperature. The scale bar represents 5 μm. Graphs represent the line profiles of the fluorescence intensity from the images on the left. (B) Percentage of colocalization calculated using 100 cells from 3 independent experiments in (A). Error bars represent SD. P-values were calculated using an unpaired student’s t-test. (C) Illustration of the imaging process in (D). Cells were treated with NaN3 and NaF for 5 min and then with R18 for 10 min while incubating with ATP inhibitors for the first observation. After removing NaN3 and NaF after a 3-min wash, the cells were observed again. (D) Colocalization between Dpm1–GFP and R18 in WT cells treated with/without NaN3 and NaF, followed by washing out the inhibitors. The scale bar represents 5 μm. Graphs indicate the line profiles of the fluorescence intensity from the images on the left. (E) Percentage of colocalization calculated using 100 cells from 3 independent experiments in (D). Error bars represent SD. P-values were calculated using one-way ANOVA. (F) Model of the P4-ATPase flippase. The ATPase forms a functional complex with the β-subunit to translocate phosphatidylserine (PS) and phosphatidylethanolamine (PE) from the exoplasmic to the cytoplasmic leaflet. There are five types of flippases, and their subunits in *S. cerevisiae* are listed. (G) R18 distribution in flippase-defective cells. *drs2*Δ cells exhibit a dot pattern (marked by arrows in the image), *dnf1*Δ *dnf2*Δ and *dnf3*Δ cells exhibit an ER pattern, and the suppression of Neo1 shows a clear circle pattern. The scale bar represents 5 μm. The phenotypes of the yeast cells representing each pattern are illustrated to the right. Graphs indicate the line profiles of the fluorescence intensity in the images on the left. (H) R18 distribution in β-subunit deletion cells. *cdc50*Δ cells exhibit a dot pattern (marked by arrows in the image) that phenocopies the pattern observed in *drs2*Δ cells. *lem3*Δ and *crf1*Δ cells exhibit an ER pattern that phenocopies the pattern observed in *dnf1*Δ *dnf2*Δ and *dnf3*Δ cells. The phenotypes of the yeast cells representing each pattern are illustrated to the right. The scale bar represents 5 μm. Graphs indicate the line profiles of the fluorescence intensity in the images on the left. (I) Graph representing the proportion (average ± SD) of cells with different patterns. n > 100 cells in 3 independent experiments. (J) Autophagic activity of flippase-defective cells measured using the GFP–Atg8 processing assay. WT and flippase-defective cells carrying a GFP–Atg8 plasmid were treated with 200 ng/mL rapamycin for 2 h to induce autophagy. Western blot analysis was performed using anti-GFP antibodies. The upper bands represent GFP-tagged Atg8 (GFP-Atg8), and the lower bands represent cleaved GFP (free GFP). (K) Quantification of the free-GFP / GFP–Atg8 ratio (mean ± SD, from three independent experiments). (L) The same samples are used in (J). Western blot analysis was performed using an anti-Ape1 antibody. The upper bands represent precursor Ape1 (prApe1), the middle bands represent mature Ape1 (mApe1), and the lower bands represent nonspecific signals that can be used as an internal control (*). (M) Quantification of the mApe1/total Ape1 ratio (mean ± SD, from three independent experiments).

Based on the above results, we hypothesized that flippase is involved in the internalization of R18. Flippase is a type 4 P-type ATPase (P4-ATPase) that catalyzes the translocation of phospholipids from the extracellular leaflet to the cytoplasmic leaflet^62^. Budding yeast exhibit five P4-ATPases (Drs2, Dnf1, Dnf2, Dnf3, and Neo1) that form heterodimeric complexes with noncatalytic subunits of three Cdc50 family member proteins, namely, Cdc50, Lem3, and Crf1^63, 64^ (Figure 2F). Among them, Drs2, Dnf1/Dnf2, and Dnf3 form complexes with Cdc50, Lem3, and Crf1, respectively. *Neo1* is an essential gene by itself^65^. Therefore, the phenotypes observed in P4-ATPase-deficient cells should be copied to those observed in cells lacking its subunits ^63, 66^. We examined R18 localization in flippase-defective cells. ER staining was defective in *drs2*Δ cells, consistent with the pattern observed in *cdc50*Δ cells, which lack the β-subunit of the P4-ATPase flippase complex. R18 staining was restricted to the PM as discrete dots (Figure 2G and 2H). Furthermore, Neo1-defective cells exhibited a distinct circular pattern, suggesting that R18 is blocked at the PM (Figure 2G and 2H). Quantification of the different patterns revealed a significant difference, suggesting that the Drs2, Cdc50 or Neo1 plays crucial roles in the internalization of R18 (Figure 2I). Notably, Drs2, Cdc50 or Neo1 are required to establish plasma membrane phosphatidylserine (PS) / phosphatidylethanolamine (PE) asymmetry^67^. Our recent study also showed that R18 is blocked at the PM in PE-deficient yeast cells (*psd1*Δ *psd2*Δ)^68^. This defect may be caused by the loss of PE asymmetry at the PM by the defects in Drs2, Cdc50 or Neo1^67^.

Considering that R18 labels ARSs, we hypothesized that the absence of flippase would result in functional defects in autophagy. To test this hypothesis, we performed the GFP–Atg8 processing assay. A centromeric plasmid expressing GFP–Atg8 was transformed into the WT cells, *atg1*Δ cells (the negative control), and each flippase-deficient cell line. Samples were collected and analyzed by western blotting using GFP-specific antibodies. The significant increase in free-GFP levels in WT cells represents the normal progression of autophagy (Figure 2J and 2K). The processing of GFP–Atg8 in *lem3*Δ, *drs2*Δ, *cdc50*Δ, *dnf3*Δ, *crf1*Δ, and Neo1-defective cells was identical to that in the WT cells, indicating that the flippases are not essential for autophagy (Figure 2J and 2K).

We also investigated selective autophagic activity using the Ape1 maturation assay. In selective autophagy, the precursor Ape1 (prApe1) forms oligomers, which are transported from the cytoplasm into a vacuole and processed into mature Ape1 (mApe1)^38^. Both prApe1 and mApe1 were detected using western blotting. Each flippase-defective cell line, but not *atg1*Δ cells, exhibited processing ability comparable to WT cells (Figure 2L and 2M), suggesting that selective autophagic activity was normal in flippase-defective cells. These findings indicate that flippase does not affect autophagy.

### Transfer of R18 from the PM to the ER depends on the oxysterol-binding protein (OSBP)-related proteins (ORPs)

To investigate how R18 is transported from the PM to the ER, we first examined the contribution of tethers at the PM–ER MCS^69^. R18 localized to the ER in tetherΔ cells and colocalized with the ER marker Sec61–GFP (Figure S3A). We then constructed a “super tetherΔ” strain by combining *ice2*Δ with tetherΔ. Even in this background, R18 still colocalized with the ER marker Scs2–GFP (Figure 3A), indicating that the transfer of R18 from the PM to the ER does not depend on PM–ER tethers. We noticed that FM4-64 is internalized through endocytosis in 10–15 min^60^. In contrast, our results revealed a rapid transfer of R18 from the PM to the ER in <3 min (Figure 2D). This transfer time is consistent with the time taken for phospholipid transfer by LTPs at the PM–ER membrane contact sites^70, 71^. Furthermore, unlike FM4-64, which primarily stains the organelles in the endocytosis pathway, R18 stains only the ER and ARSs.

**Figure 3.**
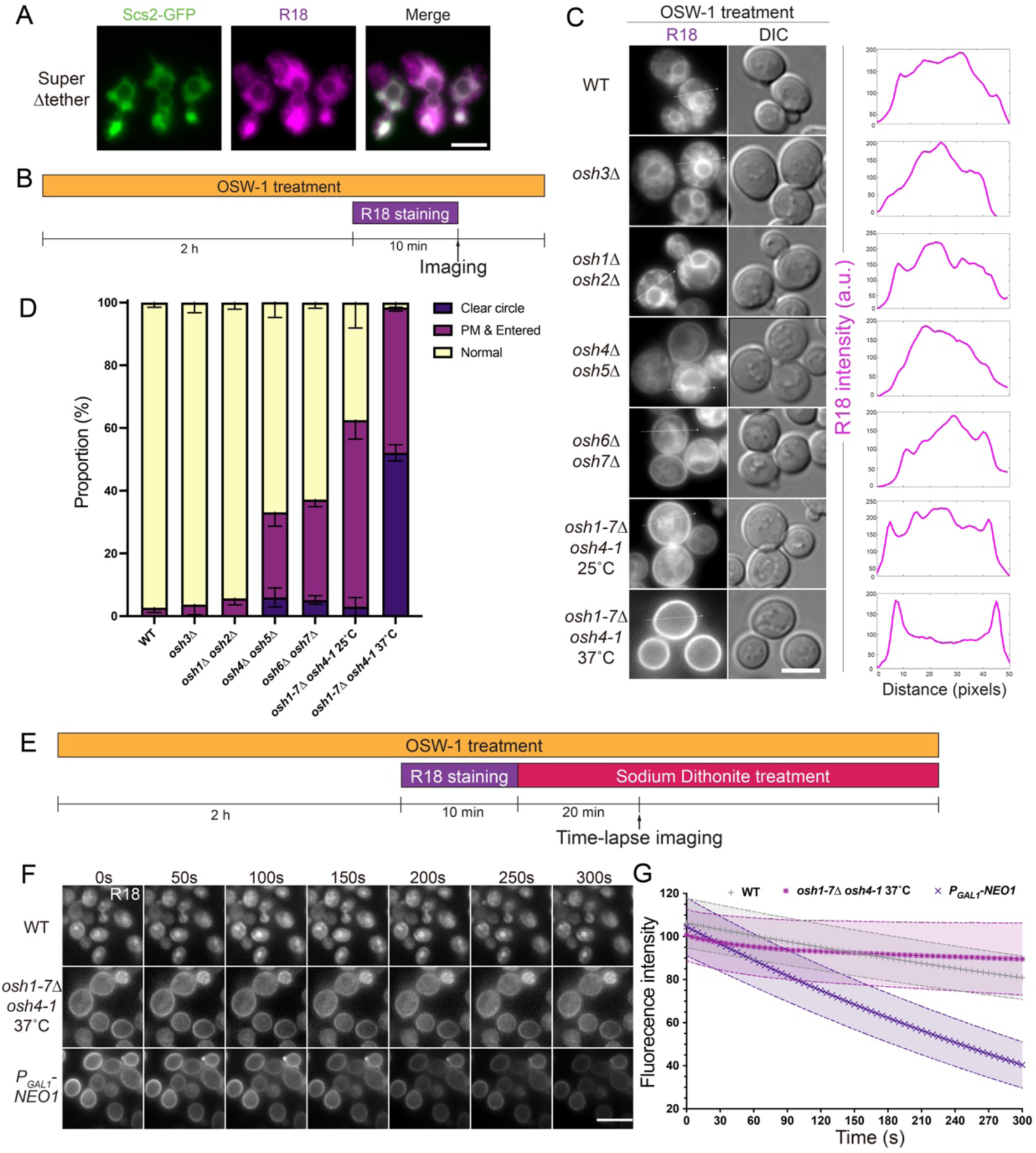
R18 transfer from the plasma membrane to the ER requires oxysterol-binding protein (OSBP)-related proteins (ORPs). (A) Colocalization between Scs2–GFP and R18 in super tetherΔ cells. The scale bar represents 5 μm. (B) The procedure for the experiment in (C). Cells were stained with R18 in the presence of 10 μM OSW-1. (C) R18 distribution in WT and Osh deletion cells. Graphs indicate the line profiles of the fluorescence intensity in the images on the left. The scale bar represents 5 μm. (D) Graph representing the proportion (average ± SD) of cells with different patterns. n > 100 cells in 3 independent experiments. (E) The procedure for the experiment in (F). Cells were stained with R18 in the presence of 10 μM OSW-1. Imaging was started 20 min after the sodium dithionite treatment. (F) Time-lapse imaging of WT, Osh deletion, and Neo1-suppressive cells. The scale bar represents 10 μm. (G) Fluorescence intensity of R18 during the time-lapse imaging in (F). Error bars represent SD; n = 7.

**Figure 4.**
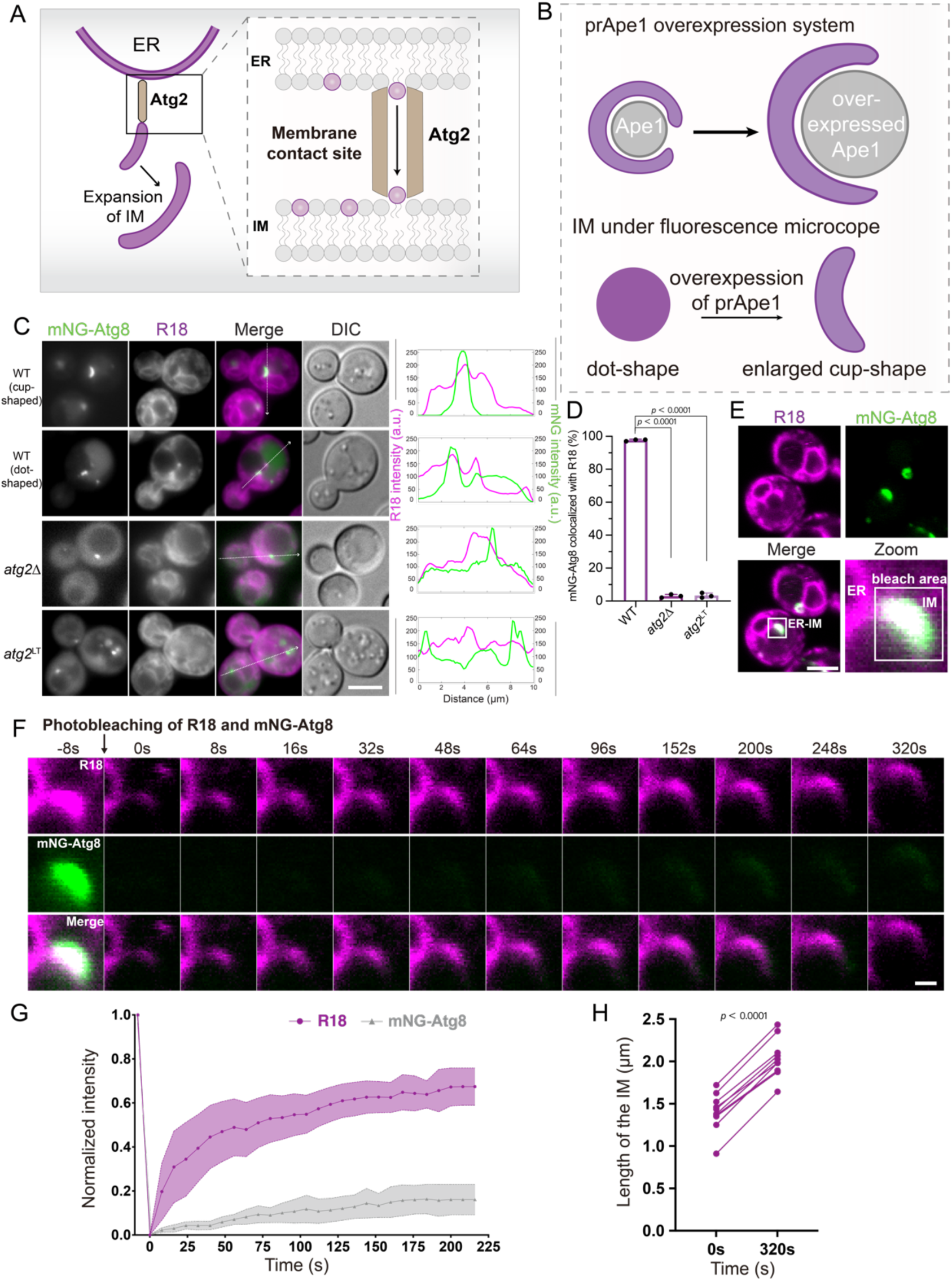
Phospholipids are transferred from the ER to the IM at the ER–IM MCS via Atg2. (A) Model of lipid transfer activity from the ER to the IM via Atg2 at the ER–IM MCS. (B) Visualization of the IM as a cup-shaped structure induced by prApe1 overexpression. (C) Colocalization between mNG-Atg8 and R18 in WT, *atg2*Δ, and *atg2*^LT^ mutant cells. The scale bar represents 5 μm. Graphs represent the line profiles of the fluorescence intensity from the images on the left. (D) Percentage of colocalization calculated using 100 cells from 3 independent experiments in (C). WT includes both dot-and cup-shaped colocalization. Error bars represent SD. P-values were calculated using one-way ANOVA. (E) IM (mNG–Atg8) is labeled with R18 in prApe1-overexpressing WT cells and visualized under the confocal microscope. The square indicates the area that was photobleached. The scale bar represents 5 μm. (F) FRAP analysis. Images were captured at the indicated time points. The scale bar represents 1 μm. (G) Fluorescence recovery of the IM (mNG-Atg8) and R18 after photobleaching. Error bars represent SD; n = 7. (H) Quantification of the length of the IM at 0 and 320 s. p-values were calculated using a two-sided paired student’s t-test.

We hypothesized that the transfer of R18 from the PM to the ER was carried out by LTPs at the PM–ER MCS.

OSBP ORPs are phospholipid transfer proteins localized to MCSs^72^. In yeast, seven homologs (Osh1–Osh7) mediate nonvesicular sterol transport at the PM–ER MCS^72^. We examined several Osh deletion cells, including *osh3*Δ, *osh1*Δ *osh2*Δ, *osh4*Δ *osh5*Δ, and *osh6*Δ *osh7*Δ. Because deletion of *OSH1-7* is lethal to cells, we used cells harboring the temperature-sensitive allele *osh4-1* in combination with deletions of the other six *OSH* genes (*osh*Δ *osh4-1*). These cells grow at the permissive temperature of 25°C but not at the nonpermissive temperature of 37°C. R18 displayed normal ER localization in WT cells, whereas no notable defects in R18 localization were observed in Osh deletion cells (Figure S3B). But R18 showed partially the PM pattern in *osh1*Δ *osh2*Δ, *osh6*Δ *osh7*Δ, and *osh*Δ *osh4-1* cells (Figure S3B). These results suggest that the transport of R18 from the PM to the ER might be impaired in these strains. Thus, we used orsaponin (OSW-1), a natural compound known to inhibit OSBP function^73, 74^, to further validate the role of OSBP in R18 transport. Treatment with OSW-1 caused a noticeable increase in the proportion of cells exhibiting partial PM-localized R18 patterns in Osh deletion cells (Figure 3B–3D). In the temperature-sensitive *osh1-7*Δ *osh4-1* cells, R18 displayed a clear circular pattern on the PM under restrictive conditions (37°C), indicating a complete block in its transport to the ER (Figure 3B–D).

Given that R18 exhibited the same clear circular pattern in Neo1-defective cells, we hypothesized that R18 would remain on the outer leaflet of the PM in Neo1-defective cells and accumulate on the inner leaflet of the PM in OSBP-defective cells. To examine the localization of R18, we treated WT, Neo1-defective, and OSBP-defective cells with sodium dithionite to quench the fluorescence of R18 exposed to the extracellular environment (Figure 3E). We observed no discernible difference in R18 intensity in the ER of WT cells. However, compared with the stable R18 intensity at the PM of OSBP-defective cells, the R18 intensity at Neo1-defective cells decreased significantly (Figure 3F and 3G). This result indicates that R18 remained on the extracellular leaflet and was not internalized in Neo1-defective cells. In contrast, the clear circle pattern on the PM observed in OSBP-deficient cells indicates that R18 accumulates on the cytoplasmic leaflet of the PM and fails to be transferred to the ER. These findings support our hypothesis that flippase and OSBP facilitate R18 transport from the PM to the ER at the PM–ER MCS. Since Neo1-or OSBP-defective cells do not transfer R18 from the PM to the ER, we included these strains as controls to assess the ER dependency of R18 transport to ARSs. In both cells, R18 remained at the plasma membrane and failed to label ARSs, whereas mNG–Atg8 puncta still formed (Figure S3C). These results suggest that R18 must reach the ER before being delivered to ARSs.

### Phospholipids are transferred from the ER to the IM at the ER–IM MCS via Atg2

Autophagosome formation requires a constant supply of phospholipids. For several decades, the source of the lipids essential for autophagosome formation has been a central and longstanding enigma in the field of autophagy. Previous *in vitro* studies suggested that Atg2 acts as an LTP^16, 18, 26, 27^. Moreover, Atg2 localizes between the ER and IM^40, 41^. Therefore, we hypothesized that Atg2 acts as an LTP at the ER–IM MCS (Figure 4A). Despite a recent electron microscopic study highlighting the importance of de novo-synthesized phosphatidylcholine in autophagosome formation^34^, direct evidence revealing lipid transfer at the ER–IM membrane contact site in living cells is still lacking, primarily due to the absence of a suitable probe to monitor phospholipid dynamics. Given that R18 stains both the ER and IM (Figure 1), we tried to use R18 as a probe to visualize phospholipid transfer from the ER to the IM *in vivo*.

The IM visualized using mNG–Atg8 appeared as a cup-shaped structure following prApe1 overexpression (Figure 4B). A recent *in silico* study has predicted an Atg2 lipid transfer mutant (*atg2*^LT^) that the density of phospholipid in the hydrophobic cavity is discontinuous^75^. We constructed the Atg2 lipid transfer mutant. In this mutant, Ape1 maturation was not detected, showing that the process of autophagy is completely blocked, whereas full length Atg2 was expressed (Figure S4A and S4B). In addition, mNG–Atg8 localized adjacent to Sec13–mRuby, suggesting that *atg2*^LT^ mutant can tether ERES and the IM (Figure S4C).

In WT cells, R18 colocalized with mNG-Atg8 as both cup-shaped structures and punctate Atg8 structures (Figure 4C and 4D). In contrast, *atg2*Δ cells showed a dot pattern without R18 as previously reported^51^. Moreover, *atg2*^LT^ cells showed the same pattern with *atg2*Δ cells (Figure 4C and 4D). One group has reported that a phospholipid transfer defective mutant of Atg2 is defective in autophagosome formation^76^. Recently, we have narrowed down the critical loop region of Atg2 and generated the *atg2*Δ21 mutant^77^. This Atg2Δ21 protein is defective in bridge-type lipid transfer activity *in vitro*^77^. R18 staining of mNG-Atg8 labeled structures is also blocked in this mutant *in vivo*^77^. These facts suggest that Atg2^LT^ protein also has a defect in the bridge-type lipid transfer activity. Collectively, these results suggest that the transfer of R18 from the ER to the IM requires the presence of a broad and continuous hydrophobic groove within the bridge-like architecture of Atg2^77^.

We next confirmed that R18 stained the IM, as observed under a confocal microscope (Figure 4E). We then performed the fluorescence recovery after photobleaching (FRAP) assay to investigate the dynamics of lipid transfer. We performed photobleaching on both R18 and mNG signals within the expanding IM and monitored their fluorescence recovery (Figure 4F, Supplemental Video 1A-C). The R18 fluorescence recovered rapidly, with a half-recovery time of 10–30 s, suggesting efficient phospholipid transfer from the ER to the IM at the ER–IM MCS (Figure 4F and 4G). Conversely, mNG–Atg8 fluorescence exhibited only limited recovery, suggesting that the supply of Atg8 to the IM during expansion is limited (Figure 4F and 4G). We assume that Atg8 localized at the IM remains essentially constant and the Atg8 pool is used for IM expansion. However, we cannot exclude the possibility that the limited supply of Atg8 is also important for IM expansion. These findings indicate that phospholipid transfer occurs from the ER to the IM at the ER–IM MCS. Furthermore, the length of the IM at 320 s was increased compared with that at 0 s (Figure 4F and 4H), indicating that the supply of Atg8 to the IM is not required during IM expansion.

### Phospholipids are transferred from the IM to the ER after autophagy termination

Although the prApe1 overexpression system enlarged the IM, it can obstruct the formation of a complete autophagosome. Therefore, we focused on the IM when it reached its maximum length during prolonged autophagy (Figure 5A). We performed photobleaching of R18 in a large region encompassing the IM, ER–IM MCS, and adjacent ER areas and subsequently monitored the fluorescence recovery at each area (Figure 5B and 5C, Supplemental Video 2A-C). The IM length did not increase after reaching the maximum length (Figure 5D). The R18 signal at the IM did not recover, indicating that phospholipid transfer from the ER to the IM had ceased (Figure 5C and 5E). We observed a normal recovery curve for R18 at the ER, showing a rapid and distinct recovery. The R18 signal at the ER–IM MCS was expected to recover at a rate comparable to that of the R18 signal at the ER. However, the fluorescence recovery was partial (Figure 5C and 5E). This outcome indicates the possible transfer of phospholipids from the quenched IM to the ER–IM MCS, indicating a previously unrecognized reversed phospholipid flow from the IM to the ER.

**Figure 5.**
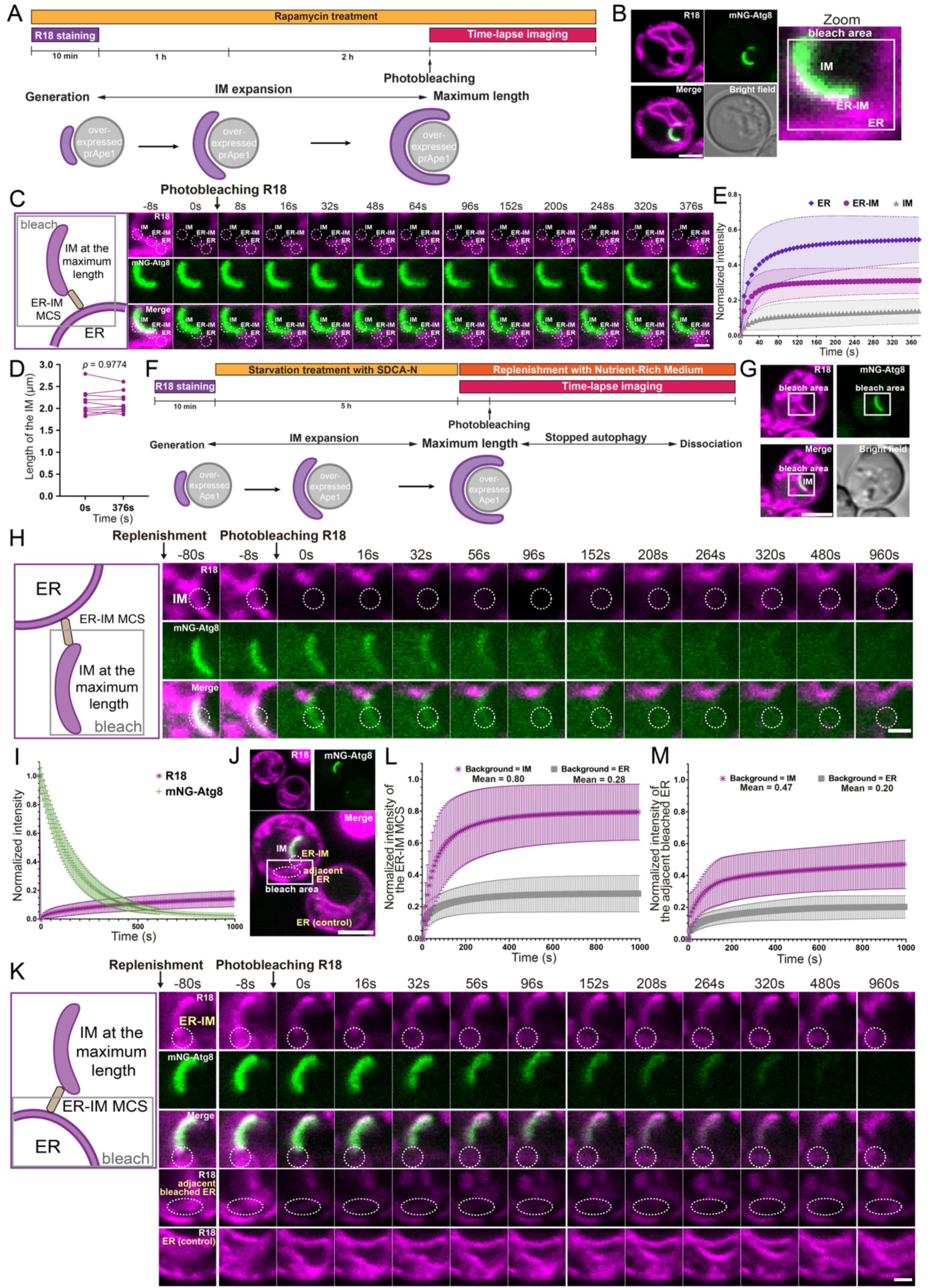
Phospholipids are transferred from the IM back to the ER after the termination of autophagy. (A) Procedure for the experiment in (C). prApe1-overexpressed cells were treated with rapamycin, and the FRAP analysis was performed as indicated. (B) The IM (mNG–Atg8) is labeled with R18 in prApe1-overexpressing WT cells under the confocal microscope. The square indicates the area that is to be photobleached in (C). The scale bar represents 2 μm. (C) Schematic drawing of the photobleached area (left). During the FRAP analysis, the fluorescence intensity of R18 at three points (ER, ER–IM, and IM) was monitored (right). Images were captured as indicated. The scale bar represents 1 μm. (D) Quantification of the length of the IM after reaching a maximum length. P-values were calculated using a two-sided paired student’s t-test. (E) Fluorescence recovery of R18 at the IM, ER–IM, and ER after photobleaching. Error bars represent SD; n = 7. (F) The procedure for the experiments in (H and K). prApe1-overexpressing cells were starved in SDCA-N medium for 5 h, and nutrient-rich medium was then added to terminate autophagy, followed by FRAP analysis as indicated. (G) IM (mNG–Atg8) is labeled with R18 in prApe1-overexpressing WT cells and visualized under a confocal microscope. The square indicates the area that was photobleached in (H). The scale bar represents 2 μm. (H) Schematic drawing of the photobleached area (left). The fluorescence intensities of mNG–Atg8 and R18 at the IM were monitored (right). Images were captured as indicated. The scale bar represents 1 μm. (I) Fluorescence recovery of mNG-Atg8 and R18 at the IM after photobleaching. Error bars represent SD; n = 7. (J) IM (mNG–Atg8) is labeled with R18 in prApe1-overexpressing WT cells and visualized under a confocal microscope. The square indicates the area that was photobleached in (K). The scale bar represents 5 μm. (K) Schematic drawing of the photobleached area (left). The fluorescence intensity of R18 at the ER–IM MCS was monitored (right). The scale bar represents 1 μm. (L) Fluorescence recovery of R18 at the ER–IM MCS, calculated using the IM and unbleached ER as background references. Mean represents the average recovery level at the final timepoint. Error bars represent SD; n = 7. (M) Fluorescence recovery of R18 at the bleached ER adjacent to the ER–IM MCS, calculated using the IM and unbleached ER as background references. Mean represents the average recovery level at the final timepoint. Error bars represent SD; n = 7.

To test the possibility of phospholipid transfer from the IM to the ER, we investigated whether this transfer occurs after the termination of autophagy. Cells were subjected to starvation for 4 h in a nitrogen-starvation medium, stained with R18, and seeded onto a glass-bottom dish. Subsequently, autophagy was terminated by replenishing the cells with a nutrient-rich medium, and the cells were examined using time-lapse microscopy (Figure 5F). We then photobleached several regions (the IM / the ER with the ER-IM MCS) to examine phospholipid transfer.

To determine whether nutrient replenishment terminates autophagy, we selectively photobleached the R18 signal and monitored the R18 (photobleached) and mNG–Atg8 (without photobleaching) signal following nutrient replenishment (Figure 5G and 5H, Supplemental Video 3A–C). We observed a marked decrease in the mNG–Atg8 signal (Figure 5G–5I), indicating the dissociation of Atg8 from the IM. Based on this result, we conclude that nutrient repletion terminates autophagosome formation. Next, we photobleached the R18 signal on the IM 80 s after nutrient replenishment (Figure 5H). We observed minimal fluorescence recovery at the IM (Figure 5G and 5H), suggesting that phospholipid transfer from the ER to the IM ceases upon autophagy termination (Figure 5H and 5I). Therefore, the cessation of anterograde lipid transfer is likely accompanied by the termination of autophagosome formation.

Next, we photobleached the ER region and monitored R18 signal recovery at the ER–IM MCS (Figure 5J and 5K, Supplemental Video 4). Time-lapse imaging revealed gradual dissociation of mNG–Atg8 at the IM after autophagy termination (Figure 5K). Following photobleaching in the ER region, R18 fluorescence was recovered at the ER–IM MCS (Figure 5K). To determine whether the phospholipids at the ER–IM MCS originated from the IM or ER, we generated a fluorescence intensity recovery curve at the ER–IM MCS, normalizing it to the intensity of the unbleached IM or ER region as the background. We observed a significant recovery of R18 fluorescence at the ER–IM MCS when the R18 fluorescence at the ER–IM MCS was normalized to that of the unbleached IM (Figure 5L), suggesting that a portion of the phospholipids in the ER–IM MCS were supplied from the IM following nutrient replenishment. Importantly, consistent with the reversed lipid flow, both Atg2 and Atg18 remained localized at the ER–IM MCS during the nutrient-replenishment assay. Time-lapse imaging showed that the Atg2 and Atg18 persisted at the contact site even after autophagy termination, supporting that the Atg2-Atg18 complex mediates lipid transfer from the IM to the ER during IM shrinkage (Figure S5; Supplemental Video 5).

To further confirm that the recovered fluorescence at the ER–IM MCS originates from the IM rather than the adjacent bleached ER (Figure 5J and 5K), we quantified the recovery of the adjacent bleached ER region. When normalized to unbleached ER, the adjacent bleached ER showed an average recovery 20% lower than that of the ER–IM MCS (28%). Furthermore, we also normalized the recovery to the unbleached IM, the adjacent bleached ER showed a much lower recovery (47%) compared to the ER–IM MCS (80%) (Figure 5L and 5M). These results indicate that at the time when fluorescence recovery at the ER–IM MCS is already apparent, the adjacent ER remains in a slower and weaker recovery phase, supporting that the recovered signal at the ER–IM MCS originates from the IM (Figure 5L and 5M). Taken together, our results indicate that the phospholipids on the IM may be transferred back to the ER during IM shrinkage after autophagy termination. Therefore, our findings provide evidence of reversible one-way lipid transfer at the ER–IM MCS.

To gain insights into the structural properties underlying the Atg2-mediated phospholipid transfer process, we employed the predictive capabilities of AlphaFold3. We modeled a system consisting of 12 molecules of 1-palmitoyl-2-oleoyl-glycero-3-phosphocholine (POPC), 12 molecules of 1,2-dioleoyl-sn-glycero-3-phosphoethanolamine (DOPE), four molecules of R18, and the RBG domain of Atg2 (residues 1–1346). The phospholipids and R18 molecules were aligned in a side-by-side configuration via the RBG domain of Atg2 (Figure 6A). Because of the inability to identify the mechanism governing the directionality of phospholipid transfer, this result suggests that phospholipids can diffuse in either direction. Cellular requirements may regulate the direction of phospholipid transfer at the ER–IM MCS. For instance, phospholipids are transferred from the ER to the IM via Atg2 during autophagosome formation. Conversely, upon autophagy termination, the phospholipid flow appears to be reversed, with the retrieval of phospholipids from the IM to the ER. The present study posits that the ER–IM MCS is a pivotal hub for regulating lipid flux between the IM and ER.

**Figure 6.**
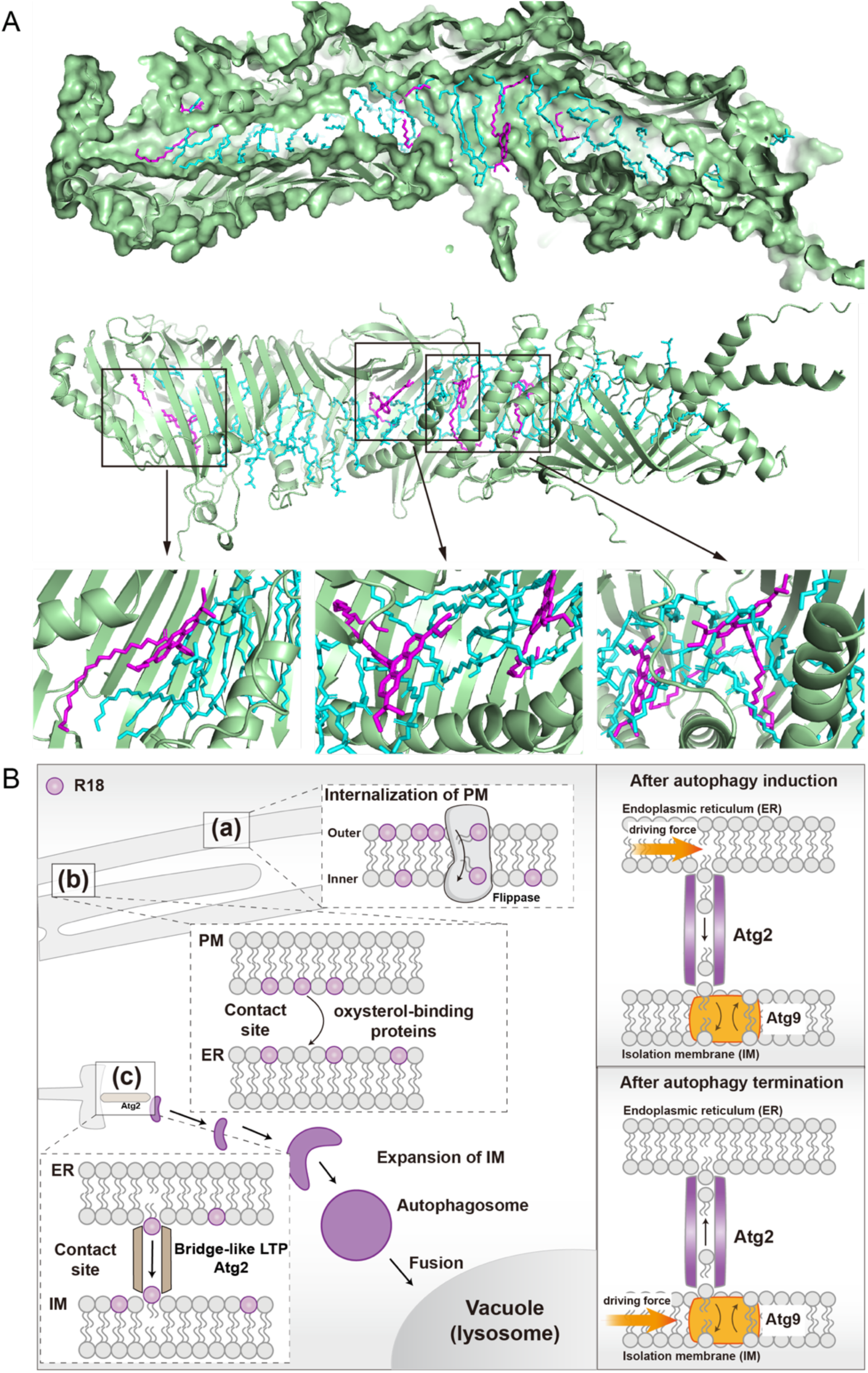
Models for R18 transport from the PM to the IM. (A) AlphaFold3 model containing Atg2 (1–1346), 12×POPC, 12×DOPE and 4×R18. Cyan represents POPC and DOPE, and magenta represents R18. (B) Models of the R18 transfer route to the IM (left). Reversible one-way lipid transfer model of Atg2 (right).

## Discussion

The study of MCSs substantially enhanced our understanding of phospholipid transfer, revealing that nonvesicular transport processes facilitate rapid and efficient lipid exchange between organelles. These processes are critical for maintaining cellular homeostasis, especially under stress or nutrient-deprivation conditions. In particular, the recent discovery of bridge-like LTPs has further significantly advanced our understanding of the structural and functional intricacies of phospholipid transfer. Bridge-like LTPs such as Atg2 and Vps13 act as molecular conduits for lipid transport, facilitating the coordinated transfer of lipids across MCSs. Despite significant structural and biochemical research progress, our understanding of lipid dynamics in living cells remains limited. Most of our current knowledge is based on *in vitro* experiments, raising concerns about the translatability of *in vitro* findings in fundamental biological research^78^.

A major constraint in our *in vivo* investigation of lipid transfer thus far has been the lack of a probe to monitor phospholipid movement. R18 was initially used to study membrane fusion during viral infection *in vitro* around 1980, coinciding with the establishment of *in vitro* assays. However, this application was subsequently abandoned by approximately 2000^50^. Our group had previously reported that R18 stains the ER^51^. The ER forms various MCSs with other organelles to facilitate lipid exchange via bridge-like LTPs. Among these LTPs, Vps13 localizes to multiple organelles and MCSs, including the ER-mitochondrion and ER-endosome MCSs^23^. R18 specifically stained the ER and occasionally labeled the vacuolar membrane, possibly due to autophagosome–vacuole fusion, but did not stain other organelles (Figure S2). Moreover, our findings indicate that R18 is specifically transferred via Atg2 at the ER–IM MCS (Figure 1 and 4). Our parallel study also confirmed Atg2 is indeed a bridge-like LTP and the transfer of R18 depends on this bridge-type lipid transfer activity at the ER-IM MCS^77^. Therefore, the transfer of R18 mediated by Vps13 and other bridge-like LTPs may be limited due to differences in the structural length and width of their hydrophobic grooves. The regulation of phospholipid transfer in various MCSs may involve a gating function restricting phospholipid transfer.

Our study analyzed phospholipid transfer pathways using R18 from the external environment of cells. In comparison with existing phospholipid traceable probes, such as FM4-64, NBD-PC, and NBD-PE, which are transported via the endocytosis pathway, we have established R18 as a novel probe for tracking lipid transfer through the nonvesicular pathway in living cells. R18 was initially internalized at the plasma membrane in a flippase-dependent manner involving Neo1 and Drs2 (Figure 2). Flippases are a type of phospholipid transporter responsible for maintaining lipid asymmetry^64^. R18 exhibited a distinct circular pattern on the PM in Neo1-defective cells, in contrast to a dot pattern in Drs2 deletion cells, suggesting the presence of entry regions or tethering sites for phospholipid translocation. Neo1 is likely required for R18 to access these entry regions on the PM. However, Neo1 is believed to be involved in recycling processes at the endosome and trans-Golgi network while also playing a role in regulating the PS/PE asymmetry of the PM^67, 79^. Most recently, we further determined that R18 internalization strictly depends on PE in the PM^68^. Our findings provide valuable insights that could inform future mechanistic studies of Neo1. We found no direct relationship between phospholipid flipping and autophagic activity, suggesting that R18 translocation occurs independently of flippase function, even when R18 reaches each ARS.

Following the treatment of *osh*Δ *osh4-1* cells with OSW-1, an OSBP inhibitor, at a restrictive temperature, R18 displayed a distinct circular blocking pattern (Figure 3). This result indicates the transfer of R18 from the PM to the ER via OSBP. One potential explanation for this phenomenon is that OSW-1 protein strongly inhibits Osh4-1 at the restrictive temperature. An alternative explanation is that undetermined OSBP(s) with low activity levels may exist that are effectively inhibited by OSW-1. Further elucidation of the molecular mechanisms underlying flippase and OSBP function requires more detailed structural studies. FRAP analysis of R18 can also be performed at the PM–ER MCS to investigate phospholipid transfer *in vivo*.

We demonstrated that R18 labels ARSs in both yeast and mammalian cells (Figure 1). In mammalian cells, the IM is sandwiched between the ER membranes, making it difficult to solely identify the flow of phospholipids within the IM^28^. Therefore, we considered *Saccharomyces cerevisiae* a suitable organism for determining the directions of phospholipid transfer at the ER–IM MCS. A major point of contention in the field of autophagy is the primary phospholipid source for autophagosome formation^80^. Our *in vivo* results strongly support the ER as the main membrane source of phospholipids, which are transferred via Atg2 to the expanding IM (Figure 4).

To investigate reversible phospholipid transfer, we leveraged the inherent defect of the prApe1 overexpression system as a key tool to enlarge the IM and prevent its closure into a complete autophagosome. We developed a nutrient-rich medium replenishment assay to terminate autophagy and focused on the IM with the maximum length (Figure 5). Atg8 dissociation may lead to IM shrinkage since we found gradual dissociation of mNG–Atg8 from the IM and reversed phospholipid flow from the IM to the ER after autophagy termination. Our FRAP data also showed that after the initial dissociation of Atg8, the IM membrane undergoes a relatively slow disassembly process suggesting that Atg8 maybe the trigger for IM shrinkage. Further research employing additional marker proteins for the IM (Atg1 and Atg16-5-12) is essential to comprehensively understand the disintegration process. We also developed a phospholipid transfer model of Atg2 (residues 1–1346) using AlphaFold3^81^ (Figure 6A). The side-by-side alignments in the RBG cavities further support our *in vivo* evidence for the reversible one-way lipid transfer via Atg2 at the ER–IM MCS. This finding highlights the critical role of bridge-like LTPs at MCSs in response to dynamic environmental conditions.

Our *in vivo* evidence demonstrated that reversible phospholipid transfer by Atg2 is a key mechanism by which cells regulate lipid homeostasis during autophagy induction and termination (Figures. 4–6). These findings also raise intriguing questions, such as how cells regulate millions of phospholipid molecules within minutes, transferring them via bridge-like LTPs at MCSs in response to environmental changes. A major issue not addressed in this study is the driving force for the reversible one-way lipid transfer at the ER–IM MCS via Atg2. Based on our findings, we hypothesized that a driving force acts on donor and receptor organelles. Furthermore, based on the studies of VPS13 in mammalian cells and the role of the Atg2–Atg9 complex in autophagy, we propose that cells regulate phospholipid transfer through a gating function in response to environmental changes. Bridge-like LTPs are likely regulated by other proteins, such as scramblase, localized at both organelles at MCSs. Nevertheless, this study successfully identified a traceable probe for phospholipids undergoing nonvesicular transport and demonstrated reversible, one-way lipid transfer via Atg2 at the ER–IM MCS. This major advancement contributes to our understanding of phospholipid transfer via bridge-like LTPs, which play a role in various cellular functions *in vivo*. Additionally, it provides insights into the chemical and biophysical aspects necessary for developing traceable probes for nonvesicular transport.

## Methods

### Yeast strains and media

*The S. cerevisiae* strains and plasmids used in this study are listed in Supplementary Tables 1 and 2, respectively. Cells were cultured in YPD (1% Bacto^TM^ yeast extract, 2% Bacto^TM^ peptone, and 2% glucose), SDCA (0.17% Difco^TM^ yeast nitrogen base w/o amino acids and ammonium sulfate, 0.5% ammonium sulfate, 0.5% Bacto^TM^ casamino acids, and 2% glucose), or SDGA (0.17% Difco^TM^ yeast nitrogen base w/o amino acids and ammonium sulfate, 0.5% ammonium sulfate, 0.5% Bacto^TM^ casamino acids, and 2% galactose) supplemented with appropriate nutrients. Plasmid amplification was performed using *Escherichia coli* strain XL1-Blue grown in LB medium (1% Bacto^TM^ tryptone, 0.5% Bacto^TM^ yeast extract, and 1% NaCl). Ampicillin was added to the LB medium at 60 μg/mL when relevant. Yeast cells were transformed using the lithium acetate method^82^. The *atg2*^LT^ mutant cells are constructed by the CRISPR/Cas9 system provided by Ellis lab (https://benchling.com/pub/ellis-crispr-tools).

To activate the Cu^2+^-inducible *CUP1* promoter, cells were cultured for 1 day in a medium containing 250 μM CuSO_4_ prior to the experiments. Autophagy was induced by transferring cells to nitrogen-starvation SD(−N) medium (0.17% yeast nitrogen base without amino acids and ammonium sulfate, and 2% glucose) or by treating cells with 0.2 μg/mL rapamycin (Sigma Aldrich). Standard protocols were used for yeast manipulation^83^.

The yeast cells were treated with a mixture of NaF and NaN_3_ at a final concentration of 10 mM for 5 min. For galactose induction of *Neo1* using the *GAL1* promoter, cells were cultured in SDGA medium supplemented with appropriate nutrients. The following day, the cells were collected and washed three times with an SDCA medium to remove the galactose, initiating the suppression of *Neo1* expression. The cells (OD_600_ = 1.0–2.0) were then resuspended in SDCA medium and incubated for 12 h to suppress *Neo1* expression.

### Fluorescence microscopy

Cells were cultured overnight, diluted, and then cultured in SDCA medium supplemented with appropriate nutrients until reaching the log phase (∼2 × 10^7^ cells per mL; OD_600_ = ∼1). For R18 staining, cells were incubated with R18 (Invitrogen) at 5 μg/mL (from a 1 mg/mL stock dissolved in dimethyl sulfoxide) for 10 min in a nutrient-rich medium at 30 C°. The cells were then washed thrice with a fresh medium. FM4-64 staining was performed as previously described^60^.

For visualization of IM, cells were cultured in the presence of 250 μM CuSO_4_ overnight to activate the *CUP1* promoter and induce prApe1 expression. The cells were then diluted and cultured in SDCA medium containing 250 μM CuSO_4_ until they reached the log phase (∼2 × 10^7^ cells per mL; OD_600_ = ∼1). Fluorescence imaging was performed using an IX83 inverted system microscope (Olympus) equipped with a UPlanSApo100× oil-immersion lens (1.40 NA; Olympus) and a CoolSNAP HQ CCD camera (Nippon Roper). U-HGLGPS (Olympus), a mercury light source system, was used to excite fluorescent proteins. U-FGFP and U-FRFP filter sets (Olympus) were used for GFP and R18 visualization, respectively. Images were acquired using the Micro-Manager software (version 1.4.6, https://micro-manager. org). Samples were prepared on 76×26-mm glass slides (S1225; Matsunami) and 18×18-mm glass coverslips (No. 1-S; Matsunami).

### Confocal microscopic analysis of mammalian cells

ER (Calreticulin-KDEL-GFP) labeling was performed using CellLight™ ER-GFP, BacMam 2.0 (C10590; Thermo Fisher Scientific). To induce autophagy, MEFs stably expressing GFP-LC3 were cultured in Earle’s Balanced Salts (E2888-500ML; Sigma) for 3 h.

Live cells seeded in 35-mm glass-bottom dishes (P35G-1.5-14-C; Mattek) were maintained at 37 °C with 5% CO_2_ and observed under a Ti2 inverted microscope (Nikon) equipped with a Nikon A1 HD25 (Nikon) confocal system and Plan Apo λ 60x oil-immersion lens (1.4 NA; Nikon). The pinhole size was set to 1.0 Airy units. For 3D imaging, z-stack images were acquired using a Plan Apo λ 60x oil-immersion lens with Galvano scanning using two frame averages, and the pinhole size was set to 0.8 Airy units. The images were captured using NIS-Elements (Nikon). The images of deconvolution were processed using Nikon’s original ER deconvolution algorithm.

### Quick-freezing and freeze-fracture replica labeling

MEF cells were starved for > 2 hours and incubated with 50 µg/mL R18 for 10 min. Cells were detached from the substrate by a brief treatment with trypsin and ethylenediaminetetraacetic acid (EDTA) and pelleted. Copper EM grids (150 mesh, Nisshin EM) were immersed in cell pellet, and sandwiched between a flat aluminum disk (Engineering Office M. Wohlwend, Sennwald, Switzerland) and a thin copper foil (20 μm thick; Nilaco). The specimens were quick-frozen using HPM 010 high-pressure freezing machine according to the manufacturer’s instruction (Leica Microsystems, Wetzlar, Germany). The frozen specimens were transferred to the cold stage of ACE900 (Leica Microsystems, Wetzlar, Germany), fractured at −102°C under a vacuum of ∼ 1 × 10-6 mbar. Replicas were made by electron-beam evaporation in three steps: carbon (6 nm thick) at an angle of 90° to the specimen surface, platinum-carbon (2 nm thick) at an angle of 45°, and carbon (10 nm thick) at an angle of 90°. Thawed replicas were treated with 2.5% SDS in 0.1 M Tris⋅HCl (pH 8.0) at 60 °C overnight.

For the double labeling of R18 and GFP-LC3, replicas were washed with phosphate-buffered saline (PBS) containing 0.1% Triton X-100 (PBST). After blocking nonspecific sites with 3% bovine serum albumin (BSA) in PBS, the samples were incubated at 4 °C overnight with 60 μg/mL mouse anti-rhodamine B monoclonal antibody (Creative Diagnostics, NY, USA) and 10.4 μg/mL rabbit anti-GFP antibody (provided by Dr Masahiko Watanabe of Hokkaido University) in PBS containing 1% BSA. The samples were then incubated with 20-fold diluted 6 nm colloidal gold AffiniPure goat anti-mouse IgG and 12 nm colloidal gold AffiniPure goat anti-rabbit IgG (Jackson ImmnunoResearch Laboratories, PA, USA) in PBS containing 1% BSA for 60 min at 37°C. The labeled replicas were observed with JEOL JEM-1400plus EM (Tokyo, Japan).

### Confocal microscopic analysis of yeast cells

A confocal laser scanning microscope FV-3000 (Olympus) equipped with a UPLSAPO60XO oil-immersion objective lens (1.42 NA; Olympus) was used. Lasers (488 and 561 nm) were employed for linear sequential excitation, and fluorescence in the ranges of 500–540 nm and 560–600 nm was recorded for mNG and octadecyl rhodamine B (R18; Invitrogen), respectively, using a resonant scanner. Samples were prepared on 76×26-mm glass slides (S1225; Matsunami) and 18×18-mm glass coverslips (No. 1-S; Matsunami). The images were analyzed using ImageJ (https://fiji.sc/).

### Quenching assay

Yeast cells immobilized on concanavalin A-coated glass-bottom dishes (Mattek) were imaged. They were then treated with 500 μL of 1M sodium dithionite dissolved in SDCA medium while being imaged under a microscope.

### SDS–PAGE and western blotting

Cells cultured in SDCA medium (2 × 10^7^ cells per mL) were subjected to lysis using alkaline trichloroacetic acid, followed by SDS–PAGE and western blotting^84^. To analyze Ape1 maturation, expression of Atg2, and GFP–Atg8 processing, SDS–PAGE was performed using 10% acrylamide gels. Proteins were transferred to polyvinylidene fluoride membranes (Immobilon-P; Millipore) using a semi-dry transfer apparatus (Trans-Blot Turbo Transfer System; Bio-Rad) at 2.5 A and 25 V for 15 min. Following the transfer, the membranes were incubated with 2% skim milk in Tris-buffered saline containing 0.05% Tween 20 (TBST) for 30 min at room temperature to block nonspecific binding. The membranes were incubated with primary anti-Ape1 (1:10,000), anti-Atg 2 (1:5000), and anti-GFP (1:5000) antibodies for 60 min at room temperature. The membranes were washed thrice with TBST and treated with horseradish peroxidase-labeled anti-rabbit secondary antibody (Promega) at a dilution of 1:5000 for 30 min, followed by an additional wash in TBST. Chemiluminescent signals were generated using an enhanced chemiluminescence reagent (GE Healthcare) and ImmunoStar LD (Fujifilm) then detected using an IR-LAS 1000 imaging system (Fujifilm).

### FRAP measurements and analysis

For the FRAP experiments assessing the IM, cells treated with rapamycin or SDCA-N were seeded onto concanavalin A-coated glass-bottom dishes (Mattek) that immobilized the cells. The experiments were performed using a confocal laser scanning microscope FV-3000 (Olympus) equipped with a UPLSAPO60XO oil-immersion objective lens (1.42 NA; Olympus). For mNG and R18 fluorescence imaging, excitation was performed using a 488 nm laser, and fluorescence was recorded at 561 nm in a linear sequential mode. Photobleaching was performed using 488 and 561 nm laser pulses. The images were analyzed using FIJI v.1.52e31. For the kinetic analysis, the relative fluorescence intensity was plotted against time, with the intensity before quenching set to 1.0 and the minimum intensity after quenching to 0.0. The data were then fitted to an exponential recovery curve: *F = A_0_(1 − exp(k_off_t))*, where *t* represents time in seconds, and A_0_ is the maximum recovery at *t* = infinity.

### Repletion assay using nutrient-rich medium

The cells were grown to the mid-log phase and then shifted to SDCA-N medium (0.17% Difco^TM^ yeast nitrogen base w/o amino acids and ammonium sulfate, 0.5% Bacto^TM^ casamino acids, and 2% glucose) to induce autophagy. The cells were then seeded on concanavalin A-coated glass-bottom dishes (Mattek) to immobilize them and were imaged. Prior to photobleaching, 1 mL of SDCA was added to the glass-bottom dish to terminate autophagy. Photobleaching and time-lapse imaging were then performed.

### Molecular modeling using AlphaFold3

Structural prediction of 4 molecules of R18, 12 molecules of DOPE and 12 molecules of POPC bound to Atg2 (residues 1–1346) was performed using AlphaFold3 v3.0.0 installed on a local computer^81^. Structural figures were prepared using PyMOL (http://www.pymol.org/pymol)^85^. A script file to run AlphaFold3 (run_1-1346_12popc_12dope_4r18.sh) and an input JSON file (1-1346_12POPC_12DOPE_4R18.json) are provided as Supplemental Materials.

## Statistical analysis

The length of the isolation membrane (IM) was quantified using the segmented line tool in ImageJ, by tracing the contour from one edge of the IM to the opposite edge. The statistical analyses of all experiments were performed using GraphPad Prism software (version 9.5.1).

## Summary of Supplemental Materials

Figures S1-S5

Supplemental Video 1-5

Supplemental text 1-2

## Acknowledgments

We thank Drs. Christopher. T. Beh, Hitoshi Nakatogawa, Jasper Rine, Ryouichi Fukuda, Kishimoto Takuma, and Kazuma Tanaka for providing plasmids and strains; Dr. Kyoka Sasaki for technical support; Dr. Yoshikazu Ohya for critical advice. We thank Dr. Kazuaki Matoba for performing the lipid transfer assay of the Atg2^LT^ protein. We thank Dr. Masahiko Watanabe for GFP-antibody and Ayaka Saito for assistance with EM sample preparation. We thank the Open Facility, Global Research Facility Alliance Center, Office for Integrated Technical Core Hub, Hokkaido University for allowing us to conduct the analysis of Replica using JEM-1400plus TEM.

This work was supported in part by JSPS KAKENHI Grant Numbers JP19H05707, JP23K20044, JP23K06667, JP24H00060 (to N.N.N.), JP 20H05313, JP22H02569 (to K.S.), and CREST, Japan Science and Technology Agency Grant number JPMJCR20E3 (to K.S., N.N.N.).

## Author contributions

K.S. supervised the project. L.H. and K.S. conceptualized the study and designed the experiments. L.H. performed the experiments and data analysis. T.M. initially visualized the colocalization of R18 and ARSs in yeast cells, and L.H. reproduced and validated these findings. Y.O., Y.H., and N.N.N. imaged the R18 distribution in mammalian cells. T.T. performed the freeze-fracture replica double labeling. T.K constructed the super tetherΔ mutant. H.L. provided insights into R18 transport by Osh proteins, and L.H. investigated it. L.H. wrote the original manuscript. L.H. and K.S. revised the manuscript.

## Declaration of interests

The authors declare no competition of interests.

## Abbreviations

ARS: autophagy-related structure
Cryo-EM: cryo-electron microscope
DOPE: 1,2-dioleoyl-sn-glycero-3-phosphoethanolamine
ER: endoplasmic reticulum
ERES: ER exit site
FRAP: fluorescence recovery after photobleaching
GFP: green fluorescent protein
IM: isolation membrane
LTP: lipid transfer protein
mApe1: mature Ape1
MCS: membrane contact site
MD: molecular dynamics
MEF: mouse embryonic fibroblasts
mNG: monomeric NeonGreen
ORP: oxysterol-binding protein-related protein
OSBP: oxysterol-binding protein
OSW-1: orsaponin
PAS: pre-autophagosomal structure/phagophore assembly site
PE: phosphatidylethanolamine
PI(3)P: phosphatidylinositol 3-phosphate
PM: plasma membrane
POPC: 1-palmitoyl-2-oleoyl-glycero-3-phosphocholine
prApe1: precursor Ape1
PS: phosphatidylserine
R18: octadecyl rhodamine B
RBG: repeating β-groove
WT: wild-type.

## Supplemental Information

**Figure S1.**
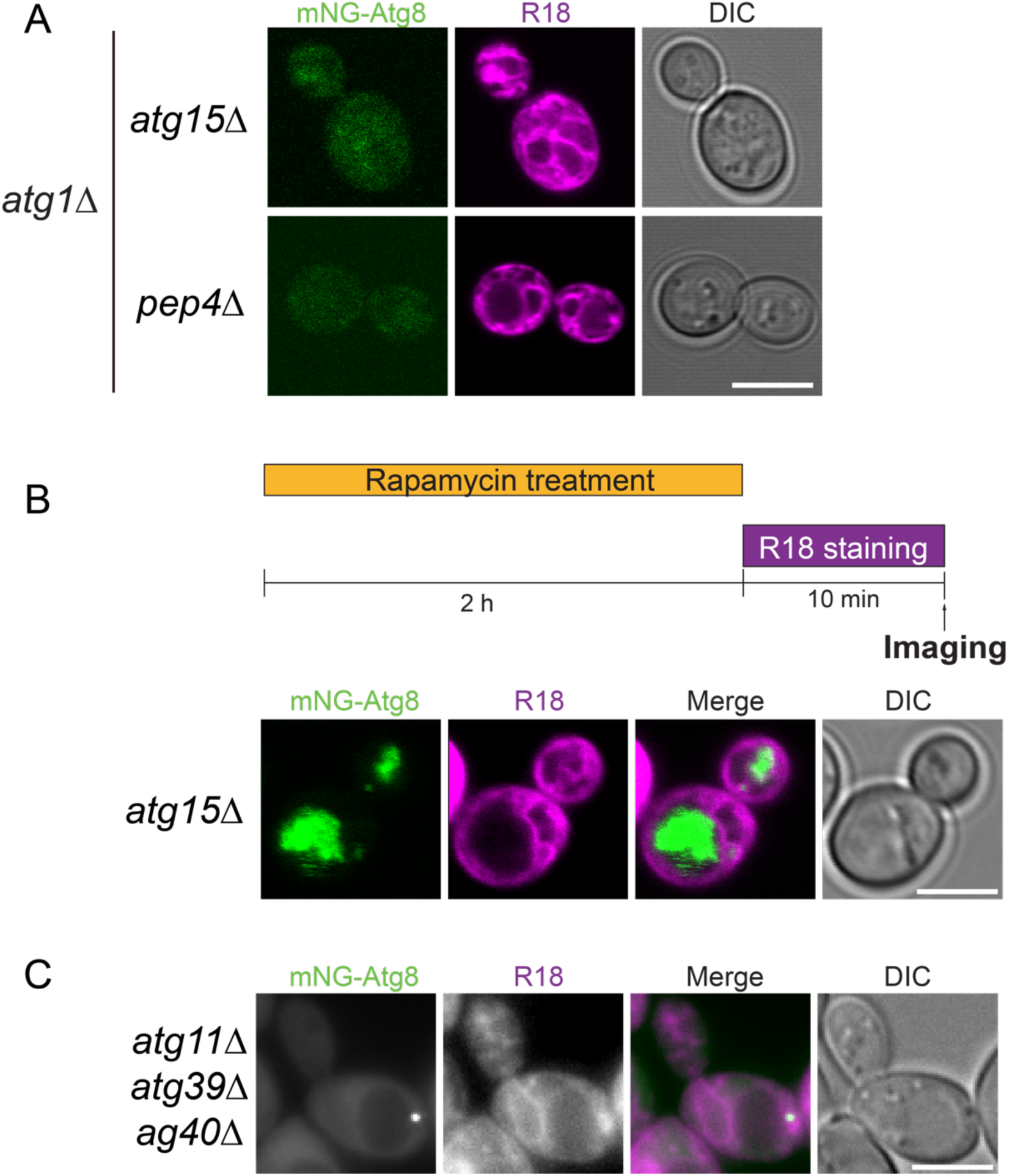
R18 staining ARSs depends on autophagy (A) mNG–Atg8 and R18 staining in *atg1*Δ *atg15*Δ and *atg1*Δ *pep4*Δ cells. Cells were stained with R18 for 10 minutes, washed with fresh medium, and then treated with 200 ng/mL rapamycin for 1 hour before imaging. The scale bar represents 5 μm. (B) Schematic of rapamycin treatment and R18 staining. Colocalization of mNG–Atg8 and R18 in *atg15*Δ cells following indicated treatment. The scale bar represents 5 μm. (C) R18 labeling in *atg11*Δ *atg39*Δ *atg40*Δ cells. Cells were stained with R18 for 10 minutes, washed with fresh medium, and then treated with 200 ng/mL rapamycin for 1 hour before imaging. The scale bar represents 5 μm.

**Figure S2.**
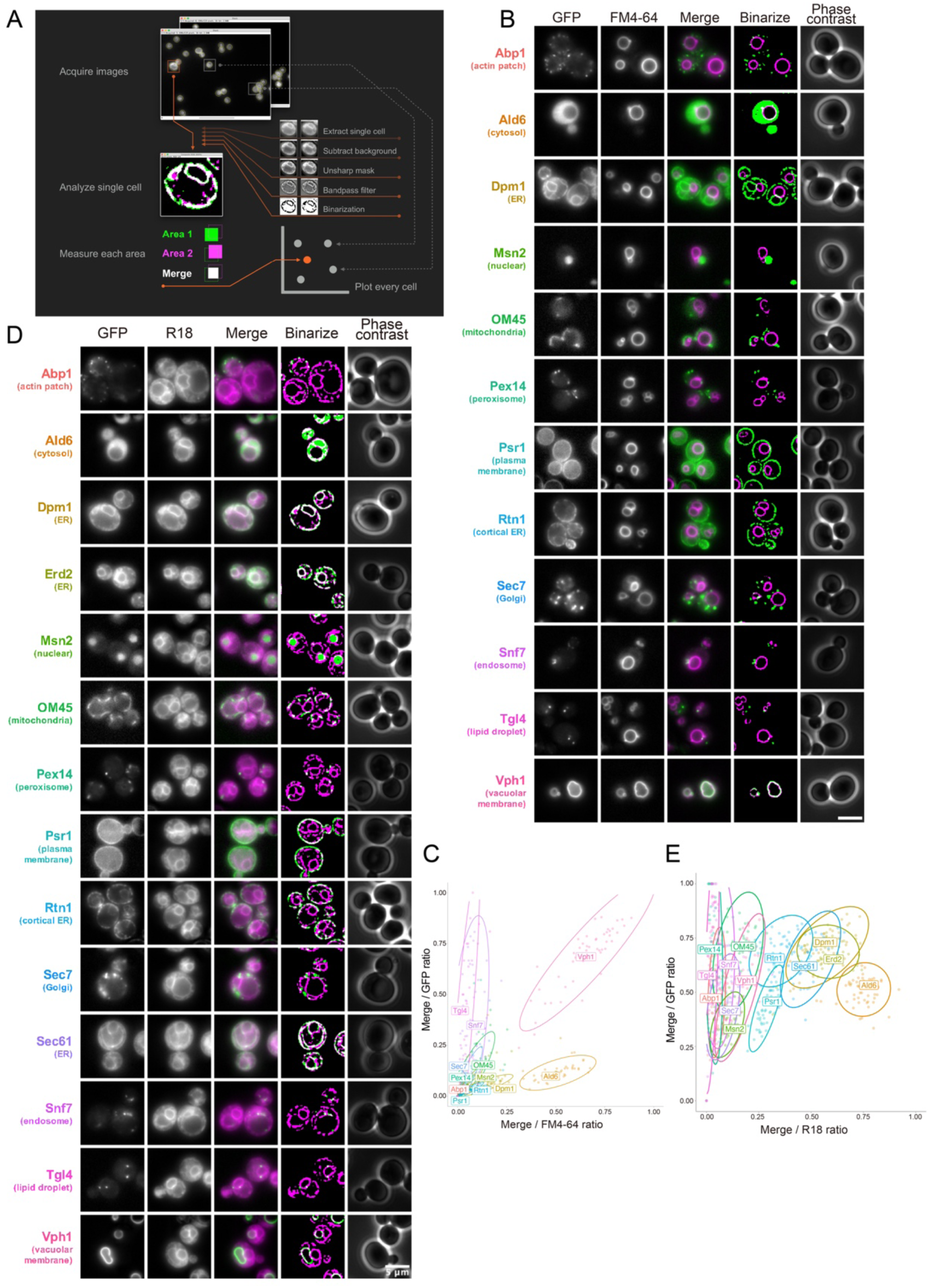
Localization analysis of FM4-64 and R18 (A) Single-cell images were extracted from the acquired images. Background brightness was subtracted using the rolling ball algorithm. Images were sharpened using unsharp masking and filtered with a bandpass filter to detect drastic intensity changes. Each structure was extracted through binarization and subsequently merged. Three areas—each fluorescence-labeled area and merged area—were measured. (B) Cells expressing GFP-tagged organelle marker proteins were grown in SDCA medium until they reached the mid-log phase and were stained with 8 μM FM4-64. The scale bar represents 5 μm. (C) Scatter plot of the merge/FM4-64 ratio (x-axis) versus the merge/GFP ratio (y-axis) with a 95% probability ellipse. At least 50 cells were analyzed for each strain. (D) Cell expressing GFP-labeled organelle marker proteins were grown in SDCA medium until they reached the mid-log phase and were stained with 10 μg/mL of R18. (E) Scatter plot of the merge/R18 ratio (x-axis) versus the merge/GFP ratio (y-axis) with a 95% probability ellipse. At least 50 cells were analyzed for each strain. The scale bar represents 5 μm.

**Figure S3.**
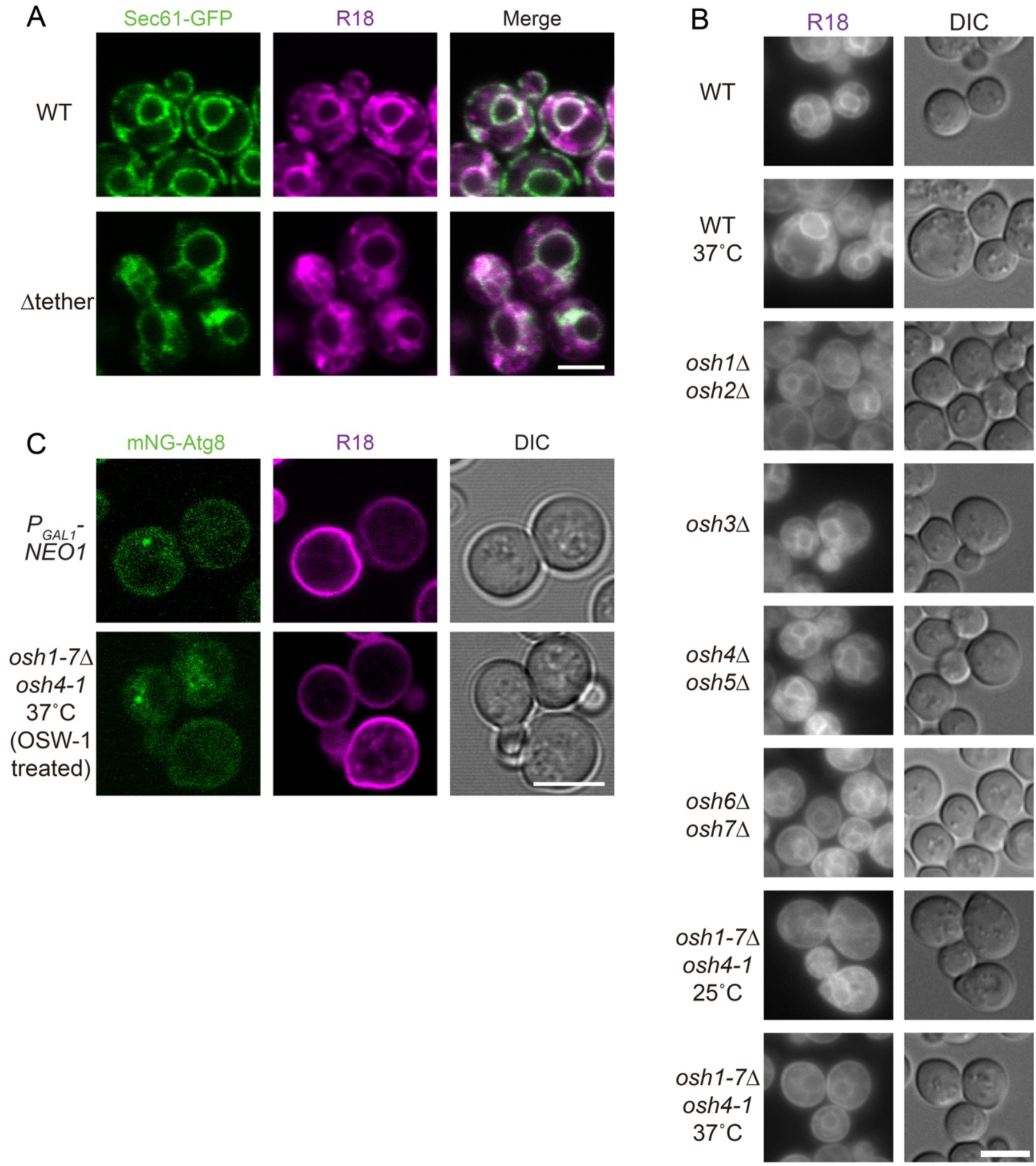
R18 distribution in the Osh and flippase mutants (A) R18 labeling of the ER in wild-type and tetherΔ cells. The scale bar represents 5 μm. (B) Cells were grown in SDCA medium to the mid-log phase at 25°C. Then, cells indicated by “37°C” were preincubated at 37°C for 1 h. The cells were stained with R18 at the indicated temperature for 10 min and then washed with fresh medium. WT, wild-type. DIC, differential interference contrast. The scale bar represents 5 μm. (C) R18 staining in flippase-or OSBP-defective cells. NEO1-defective cells and OSBP-defective cells were stained with R18. Cells were stained with R18 for 10 minutes, washed with fresh medium, and then treated with 200 ng/mL rapamycin for 1 hour before imaging. The scale bar represents 5 μm.

**Figure S4.**
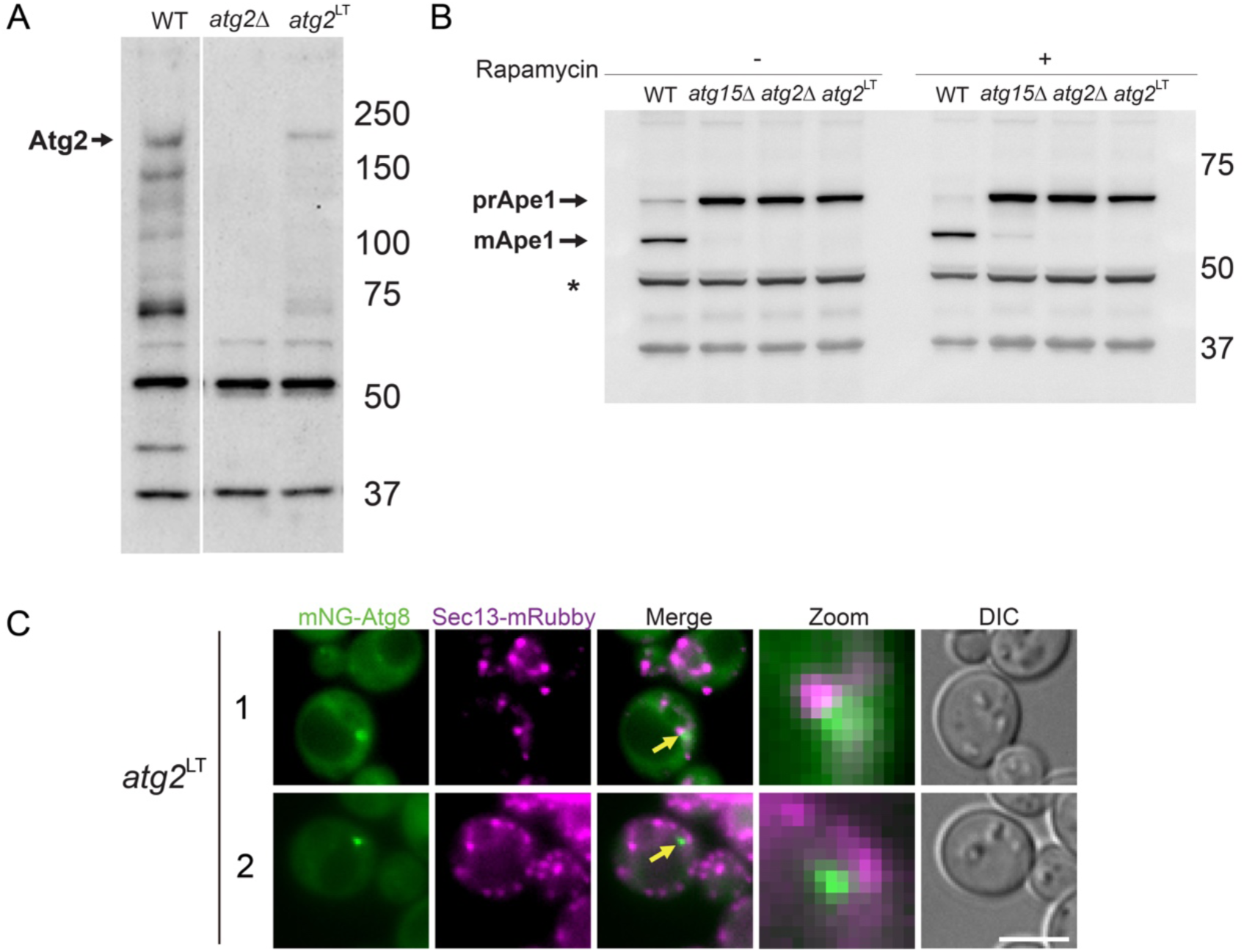
Construction of *atg2^LT^* cells. (A) Immunoblotting analysis was performed using anti-Atg2 antibodies. (B) The same samples are used in (A). Western blot analysis was performed using an anti-Ape1 antibody. The upper bands represent precursor Ape1 (prApe1), the middle bands represent mature Ape1 (mApe1), and the lower bands represent nonspecific signals that can be used as an internal control (*). (C) Colocalization between mNG-Atg8 and Sec13-mRubby in *atg2*^LT^ cells. The scale bar represents 5 μm.

**Figure S5.**
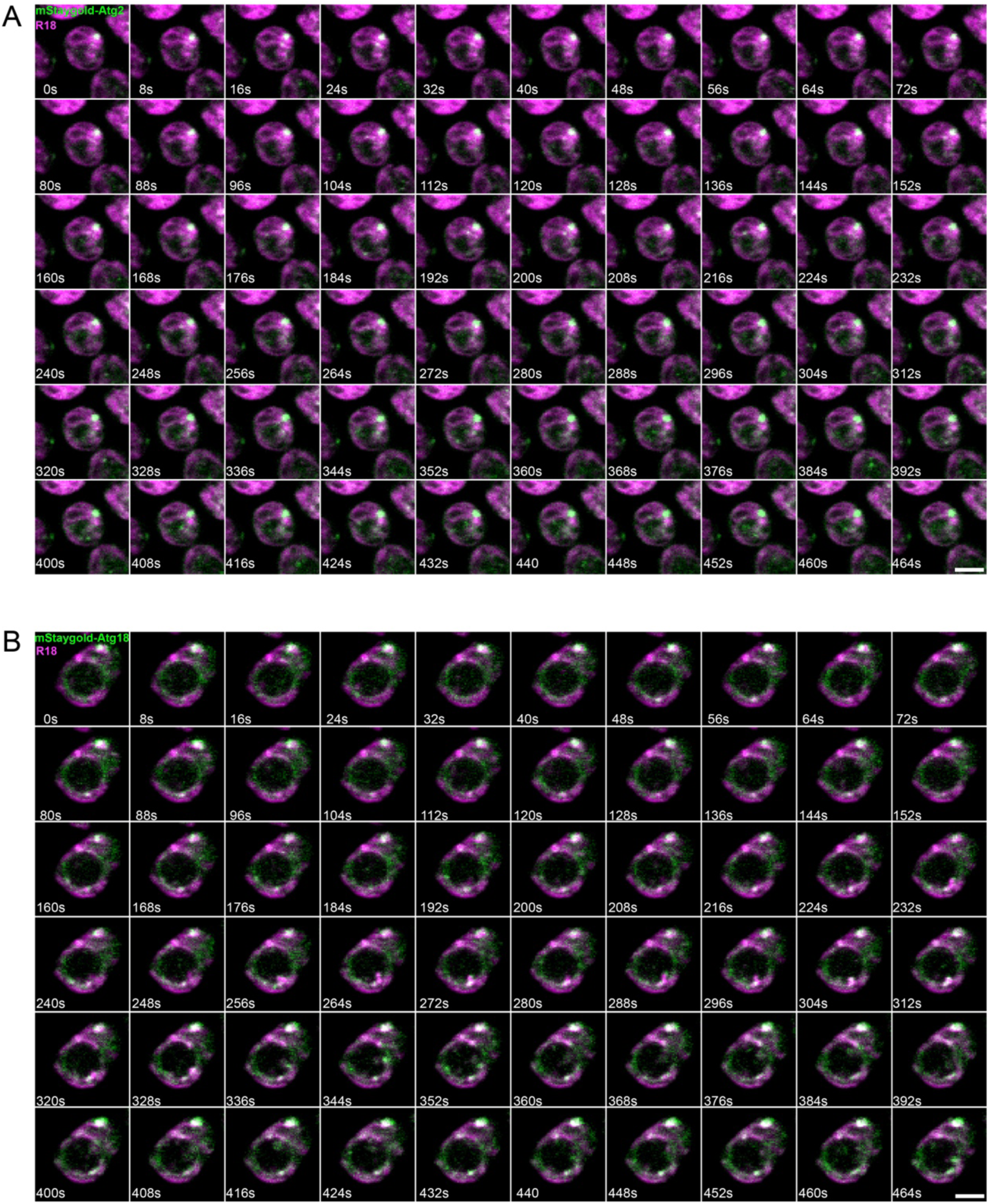
Localization of Atg2 and Atg18 during replenishment with nutrient-rich medium. (A, B) Time-lapse fluorescence microscopy of yeast cells expressing mStaygold–Atg2 (A) or mStaygold–Atg18 (B) stained with the fluorescent lipid dye R18. Autophagy was induced by nitrogen starvation for 5 h and subsequently terminated by replenishment with nutrient-rich medium, under the same conditions as in Figure 5. Numbers indicate time in seconds. The scale bar represents 2 μm.

---

**Table S1.**
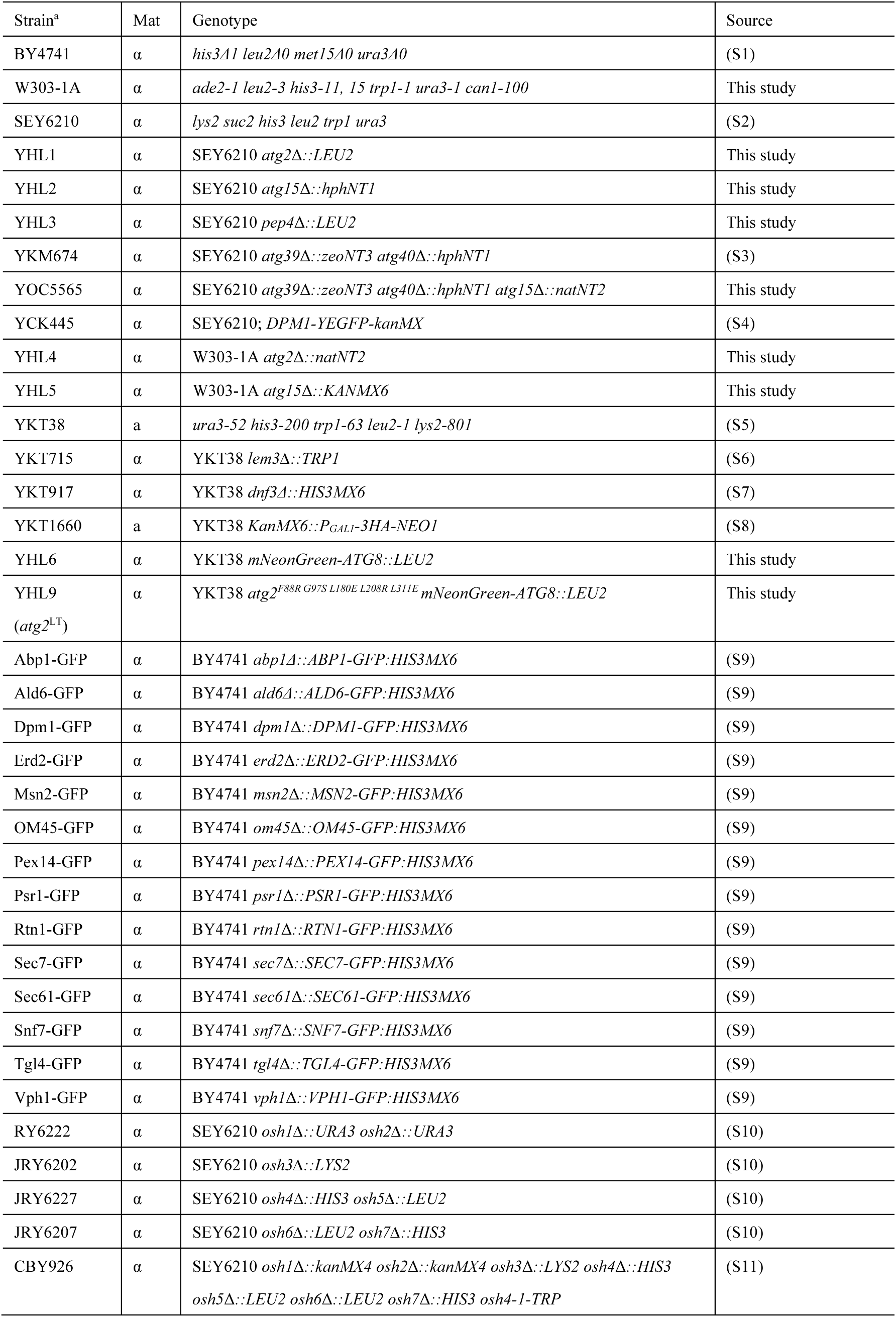
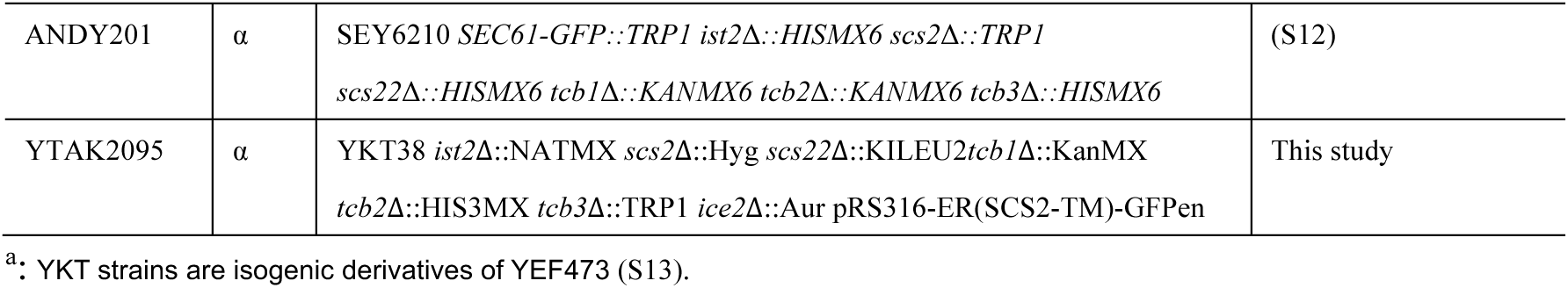
*S. cerevisiae* strains used in this study.

| Strain<sup>a</sup> | Mat | Genotype | Source |
| --- | --- | --- | --- |
| BY4741 | $\alpha$ | *his3Δ1 leu2Δ0 met15Δ0 ura3Δ0* | (S1) |
| W303-1A | $\alpha$ | *ade2-1 leu2-3 his3-11, 15 trp1-1 ura3-1 can1-100* | This study |
| SEY6210 | $\alpha$ | *lys2 suc2 his3 leu2 trp1 ura3* | (S2) |
| YHL1 | $\alpha$ | SEY6210 *atg2Δ::LEU2* | This study |
| YHL2 | $\alpha$ | SEY6210 *atg15Δ::hphNT1* | This study |
| YHL3 | $\alpha$ | SEY6210 *pep4Δ::LEU2* | This study |
| YKM674 | $\alpha$ | SEY6210 *atg39Δ::zeoNT3 atg40Δ::hphNT1* | (S3) |
| YOC5565 | $\alpha$ | SEY6210 *atg39Δ::zeoNT3 atg40Δ::hphNT1 atg15Δ::natNT2* | This study |
| YCK445 | $\alpha$ | SEY6210; *DPM1-YEGFP-kanMX* | (S4) |
| YHL4 | $\alpha$ | W303-1A *atg2Δ::natNT2* | This study |
| YHL5 | $\alpha$ | W303-1A *atg15Δ::KANMX6* | This study |
| YKT38 | a | *ura3-52 his3-200 trp1-63 leu2-1 lys2-801* | (S5) |
| YKT715 | $\alpha$ | YKT38 *lem3Δ::TRP1* | (S6) |
| YKT917 | $\alpha$ | YKT38 *dnf3Δ::HIS3MX6* | (S7) |
| YKT1660 | a | YKT38 *KanMX6::P<sub>GALI</sub>-3HA-NEO1* | (S8) |
| YHL6 | $\alpha$ | YKT38 *mNeonGreen-ATG8::LEU2* | This study |
| YHL9<br>(*atg2<sup>LT</sup>*) | $\alpha$ | YKT38 *atg2<sup>F88R G97S L180E L208R L311E</sup> mNeonGreen-ATG8::LEU2* | This study |
| Abp1-GFP | $\alpha$ | BY4741 *abp1Δ::ABP1-GFP:HIS3MX6* | (S9) |
| Ald6-GFP | $\alpha$ | BY4741 *ald6Δ::ALD6-GFP:HIS3MX6* | (S9) |
| Dpm1-GFP | $\alpha$ | BY4741 *dpm1Δ::DPM1-GFP:HIS3MX6* | (S9) |
| Erd2-GFP | $\alpha$ | BY4741 *erd2Δ::ERD2-GFP:HIS3MX6* | (S9) |
| Msn2-GFP | $\alpha$ | BY4741 *msn2Δ::MSN2-GFP:HIS3MX6* | (S9) |
| OM45-GFP | $\alpha$ | BY4741 *om45Δ::OM45-GFP:HIS3MX6* | (S9) |
| Pex14-GFP | $\alpha$ | BY4741 *pex14Δ::PEX14-GFP:HIS3MX6* | (S9) |
| Psr1-GFP | $\alpha$ | BY4741 *psr1Δ::PSR1-GFP:HIS3MX6* | (S9) |
| Rtn1-GFP | $\alpha$ | BY4741 *rtn1Δ::RTN1-GFP:HIS3MX6* | (S9) |
| Sec7-GFP | $\alpha$ | BY4741 *sec7Δ::SEC7-GFP:HIS3MX6* | (S9) |
| Sec61-GFP | $\alpha$ | BY4741 *sec61Δ::SEC61-GFP:HIS3MX6* | (S9) |
| Snf7-GFP | $\alpha$ | BY4741 *snf7Δ::SNF7-GFP:HIS3MX6* | (S9) |
| Tgl4-GFP | $\alpha$ | BY4741 *tgl4Δ::TGL4-GFP:HIS3MX6* | (S9) |
| Vph1-GFP | $\alpha$ | BY4741 *vph1Δ::VPH1-GFP:HIS3MX6* | (S9) |
| RY6222 | $\alpha$ | SEY6210 *osh1Δ::URA3 osh2Δ::URA3* | (S10) |
| JRY6202 | $\alpha$ | SEY6210 *osh3Δ::LYS2* | (S10) |
| JRY6227 | $\alpha$ | SEY6210 *osh4Δ::HIS3 osh5Δ::LEU2* | (S10) |
| JRY6207 | $\alpha$ | SEY6210 *osh6Δ::LEU2 osh7Δ::HIS3* | (S10) |
| CBY926 | $\alpha$ | SEY6210 *osh1Δ::kanMX4 osh2Δ::kanMX4 osh3Δ::LYS2 osh4Δ::HIS3*<br>*osh5Δ::LEU2 osh6Δ::LEU2 osh7Δ::HIS3 osh4-1-TRP* | (S11) |
| ANDY201 | $\alpha$ | SEY6210 *SEC61-GFP::TRP1 ist2Δ::HISMX6 scs2Δ::TRP1 scs22Δ::HISMX6 tcb1Δ::KANMX6 tcb2Δ::KANMX6 tcb3Δ::HISMX6* | (S12) |
| YTAK2095 | $\alpha$ | YKT38 *ist2Δ::NATMX scs2Δ::Hyg scs22Δ::KILEU2tcb1Δ::KanMX tcb2Δ::HIS3MX tcb3Δ::TRP1 ice2Δ::Aur pRS316-ER(SCS2-TM)-GFPen* | This study |

<sup>a</sup>: YKT strains are isogenic derivatives of YEF473 (S13).

**Table S2.**
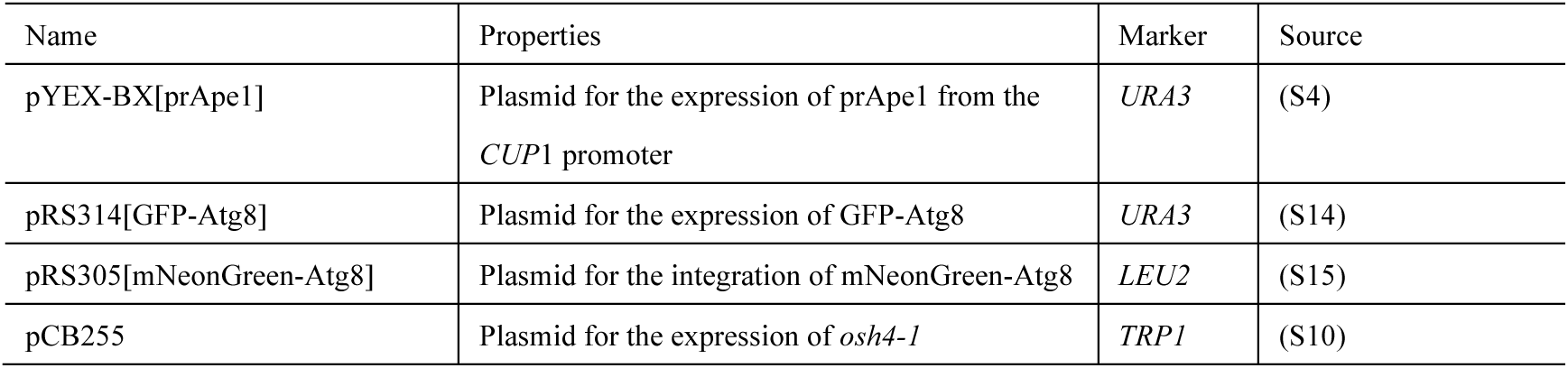
Plasmids used in this study.

## Notes

### Competing Interest Statement

The authors have declared no competing interest.

### Summary of Updates

New evidence and additional controls strengthen the idea that R18 labeling of ARSs depends on autophagy and R18 reports biologically regulated lipid transfer rather than passive diffusion (Figures 1I, 3A, S1, S3, and S4); Additional calculation evidence for reversible lipid transfer from the IM back to the ER (Figure 5); Added Atg2/Atg18 localization during nutrient replenishment (Figure S5); Cited our parallel study to collectively suggest that the transfer of R18 from the ER to the IM requires the presence of a broad and continuous hydrophobic groove within the bridge-like architecture of Atg2; We clarified that the effects of flippases and Osh proteins on R18 transfer are likely indirect and cited our work that R18 internalization depends on PE; We corrected several errors in the previous version.

